# SARS-CoV-2 impairs interferon production via NSP2-induced repression of mRNA translation

**DOI:** 10.1101/2022.01.19.476693

**Authors:** Jung-Hyun Choi, Xu Zhang, Christine Zhang, David L. Dai, Jun Luo, Reese Ladak, Qian Li, Shane Wiebe, Alex C.H. Liu, Xiaozhuo Ran, Jiaqi Yang, Parisa Naeli, Aitor Garzia, Lele Zhou, Niaz Mahmood, Qiyun Deng, Mohamed Elaish, Rongtuan Lin, Tom Hobman, Jerry Pelletier, Tommy Alain, Silvia Vidal, Thomas Duchaine, Mohammad T. Mazhab-Jafari, Xiaojuan Mao, Seyed Mehdi Jafarnejad, Nahum Sonenberg

**Author notes:** Correspondence should be addressed to N.S. and S.M.J. Lead contact: N.S. These authors contributed equally to this work (Jung-Hyun Choi, Xu Zhang, Christine Zhang).

## Abstract

Viruses evade the innate immune response by suppressing the production or activity of cytokines such as type I interferons (IFNs). Here we report the discovery of a novel mechanism by which the SARS-CoV-2 virus co-opts an intrinsic cellular machinery to suppress the production of the key immunostimulatory cytokine IFN-β. We reveal that the SARS-CoV-2 encoded Non-Structural Protein 2 (NSP2) directly interacts with the cellular GIGYF2 protein. This interaction enhances the binding of GIGYF2 to the mRNA cap-binding protein 4EHP, thereby repressing the translation of the *Ifnb1* mRNA. Depletion of GIGYF2 or 4EHP significantly enhances IFN-β production, leading to reduced viral infection. Our findings reveal a new target for rescuing the antiviral innate immune response to SARS-CoV-2 and other RNA viruses.

## Introduction

Production of type I interferons (IFN-α and IFN-β) is pivotal to antiviral immunity as a host defence mechanism^1^. Replication of SARS-CoV-2 is sensitive to type I IFN expression *in vitro*^2-4^, and the life-threatening SARS-CoV-2 infection is associated with a deficiency in type I IFN response^5,6^. In the early phase of the SARS-CoV-2 infection, a robust IFN-induced antiviral response limits viral replication and prevents severe COVID-19 illness^7,8^. Conversely, impaired production of type I IFN is associated with higher viral titers in blood and pernicious symptoms in late-stage SARS-CoV-2-infected patients^9^.

Production of type I IFNs is controlled at several levels, including transcription and translation. Notably, multiple SARS-CoV-2 proteins inhibit *Ifnb1* transcription, including NSP1, 3, 5, 6, 12, 13, 14 15, ORF3a, ORF6 and ORF7b^10,11^. Potent translational repression of *Ifnb1* mRNA is also manifested during SARS-CoV-2 infection^12^. Although SARS-CoV-2 represses general cellular mRNA translation machinery to support viral mRNA translation^12-14^, the mechanism by which it specifically represses *Ifnb1* mRNA translation is not known.

Translation of most eukaryotic mRNAs is facilitated by binding of the eukaryotic initiation factor 4E (eIF4E) to the 5′ cap structure (m7GpppN, where N is any nucleotide, and m is a methyl group). eIF4E is part of the eIF4F complex, which also contains the scaffolding protein eIF4G and the RNA helicase eIF4A^15^. Being the least abundant initiation factor, eIF4E is rate-limiting for eIF4F formation and translation initiation. The eIF4E homologous protein 4EHP (eIF4E2) also binds the cap structure but fails to initiate canonical translation because it does not bind to eIF4G. Consequently, 4EHP represses mRNAs translation upon recruitment to target mRNAs (*e*.*g*. via 4E-T protein upon recruitment by microRNAs)^16-19^. GIGYF2 [Grb10-interacting GYF (glycine, tyrosine, phenylalanine) protein 2] is another protein among others that is recruited by 4EHP to the mRNA to inhibit translation or decrease stability^20-25^. GIGYF2 participates in both 4EHP-dependent and -independent post-transcriptional repression mechanisms^20,22-24^. We recently reported the 4EHP-mediated, miR-34a-directed translational repression of *Ifnb1* mRNA^26^. This mechanism limits IFN-β production upon viral infection, likely to prohibit prolonged inflammatory responses^26^. Whether GIGYF2 is involved in the 4EHP-mediated translational repression of IFN-β is unknown.

Several large-scale proteomic studies reported the interaction of SARS-CoV-2 Non-Structural Protein 2 (NSP2) with 4EHP and GIGYF2^27-29^. Here, we document the discovery of a mechanism by which the NSP2 protein impedes IFN-β expression through translational repression of *Ifnb1* mRNA by co-opting the GIGYF2/4EHP complex, leading to evasion of cellular innate immune response and enhanced viral replication.

## Results

### NSP2 specifically interacts with GIGYF2 in the GIGYF2/4EHP translation repression complex

GIGYF2 and 4EHP play a crucial role in repressing mRNA translation via the miRNA-induced silencing complex (miRISC)^18,23,30^. We first sought to validate the interaction between NSP2 and the GIGYF2/4EHP complex in cells using the Proximity Ligation Assay (PLA). FLAG-NSP2 co-transfected with v5-GIGYF2 resulted in a strong PLA signal, which was absent in cells co-transfected with FLAG-NSP2 and v5-GIGYF1 (a paralogue of GIGYF2; Fig. 1a & b, Extended Data Fig. 1a & b). Strikingly, we did not detect any signal upon co-transfection of FLAG-NSP2 with v5-4EHP (Fig. 1a & b), which indicates that NSP2 directly interacts with GIGYF2, but not with 4EHP.

**Figure 1.**
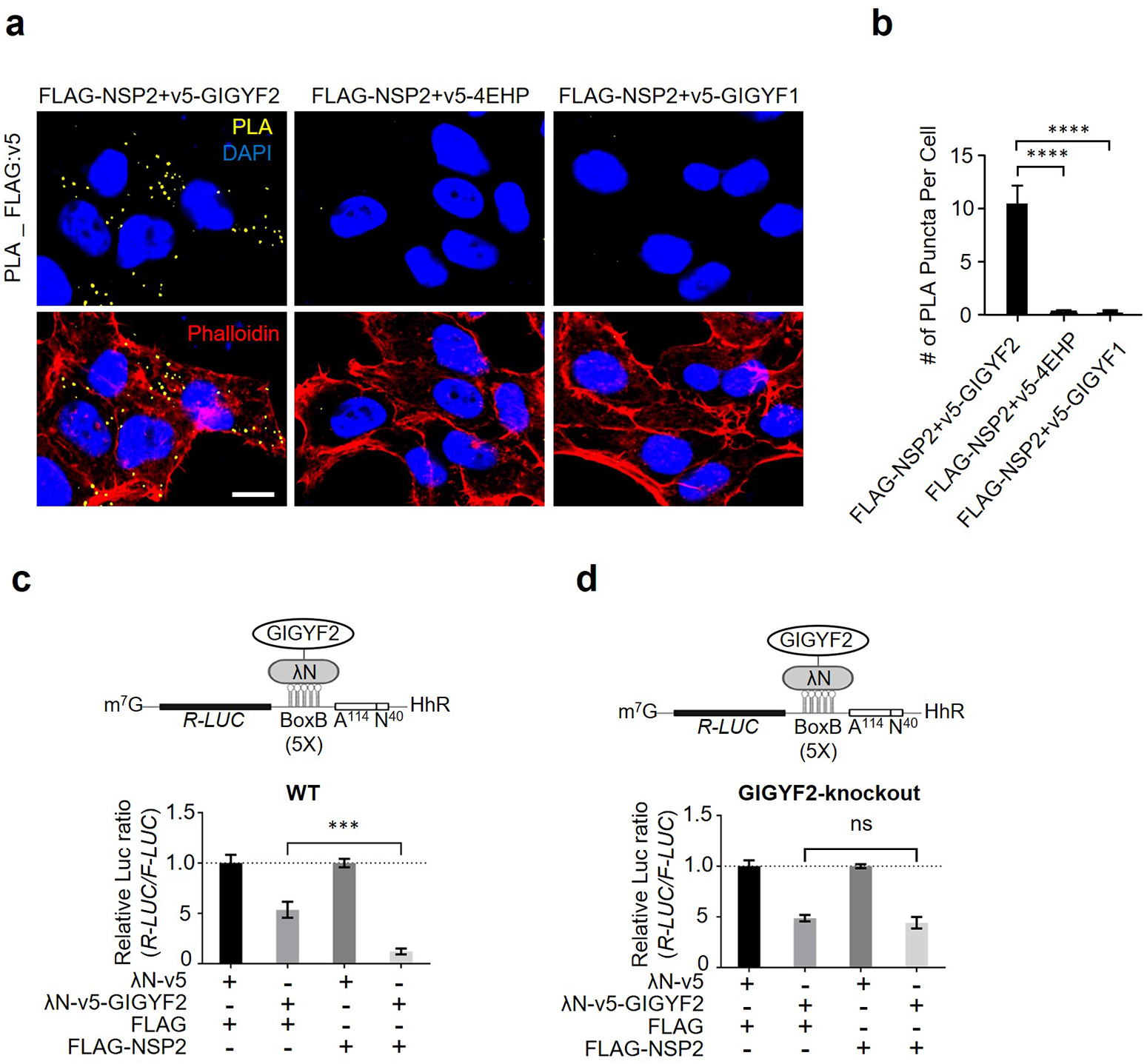
NSP2 bolsters the GIGYF2/4EHP translational repression complex. (**a**) PLA detection of NSP2-GIGYF2 interactions visible as fluorescent punctate in HEK293T cells transfected with vectors expressing v5-tagged GIGYF2, 4EHP, or GIGYF1 together with FLAG-NSP2. 24 h post-transfection cells were fixed and subjected to PLA using FLAG and v5 antibodies. PLA signals are shown in yellow. Nucleus and actin cytoskeleton were counterstained with DAPI (blue) and phalloidin (red), respectively. Scale bar= 10 µm. (**b**) Quantification of positive PLA signals in (a). The number of PLA signals from at least 30 cells was counted in each sample. ****P<0.0001 one-way ANOVA with Bonferroni’s post hoc test; n=5 independent replicates. Data are presented as mean ± SD. (**c**) WT HEK293 cells were co-transfected with plasmids expressing either λN-v5-GIGYF2 or λN-v5 as control, along with *R-Luc*-5BoxB-A114-N40-HhR and *F-Luc* (as control), followed by dual-luciferase measurement assay. Data are presented as mean ± SD (n=3). ***p<0.001, one-way ANOVA with Bonferroni’s post hoc test. The schematic shows a graphic model of λN-v5-GIGYF2 tethering system. (**d**) Analysis of λN-v5-GIGYF2 tethering-induced silencing in GIGYF2-KO cells that overexpress FLAG-NSP2. Data are presented as mean ± SD (n=3). ns= non-significant, one-way ANOVA with Bonferroni’s post hoc test. See also Extended Data Fig. 1.

Next, to investigate whether NSP2 impacts translational repression by GIGYF2, we used the λN-BoxB system to tether the λN-fused GIGYF2 to the 3′ <u>U</u>n<u>T</u>ranslated <u>R</u>egion (3′ UTR) of *Renilla* luciferase (*R-Luc*) mRNA. The reporter mRNA is protected against deadenylation by a Hammerhead ribozyme (HhR) located at its 3′ end^31,32^. We co-transfected the reporter along with FLAG-NSP2 or FLAG-empty control plasmid. While GIGYF2-tethering alone resulted in ∼50% repression [the *R-Luc*/*F-Luc* (*firefly luciferase*) ratio] compared to the counterpart λN vector, co-expression of NSP2 along with GIGYF2-tethering plasmid resulted in a stronger repression (∼88%) (Fig. 1c, Extended Fig. 1c). In contrast, *R-Luc* repression was unaffected in GIGYF2-KO cells transfected with NSP2 (Fig. 1d, Extended Fig. 1d), most probably owning to the absence of 4EHP, which is unstable in GIGYF2-depleted cells^20^. Importantly, NSP2 did not increase translational repression by tethered GIGYF1 (Extended Data Fig. 1e & f), demonstrating the specificity of NSP2-induced GIGYF2-mediated translational repression.

### NSP2 induces translational repression by bolstering GIGYF2-4EHP interaction

GIGYF2 employs both 4EHP-dependent and -independent mechanisms to translationally repress target mRNAs^20,23,33^. To study how 4EHP contributes to the GIGYF2-mediated translational repression by NSP2, GIGYF2 tethering experiments were carried out in WT, 4EHP-KO, and GIGYF2-KO cells with or without ectopic 4EHP expression. Expression of NSP2 enhanced the GIGYF2 tethering-induced silencing (from 0.41 to 0.15) in WT cells, but not in 4EHP-KO or GIGYF2-KO cells (Fig. 2a-c, Extended Data Fig. 2a-c). 4EHP expression restored GIGYF2-mediated repression (from 0.45 to 0.05 and from 0.40 to 0.08 in NSP2-overexpressing 4EHP-KO and GIGYF2-KO cells, respectively). Notably, ectopic expression of 4EHP did not affect GIGYF2-mediated repression in WT, 4EHP-KO and GIGYF2-KO cells (Fig. 2a-c). These results support the notion that NSP2 promotes GIGYF2-induced translational repression in a 4EHP-dependent manner (Fig. 2d). To test whether NSP2 bolsters the interaction of GIGYF2 and 4EHP, we carried out the PLA assay following co-transfection of v5-GIGYF2 and HA-4EHP into WT HEK293 cells with NSP2. As expected, the interaction between 4EHP and GIGYF2 was dramatically enhanced upon expression of NSP2 (FLAG: 19.8 ± 1.9, FLAG-NSP2: 61 ± 5.5 punctate per cell (Fig. 2e & f, Extended Data Fig. 3 a-c).

**Figure 2.**
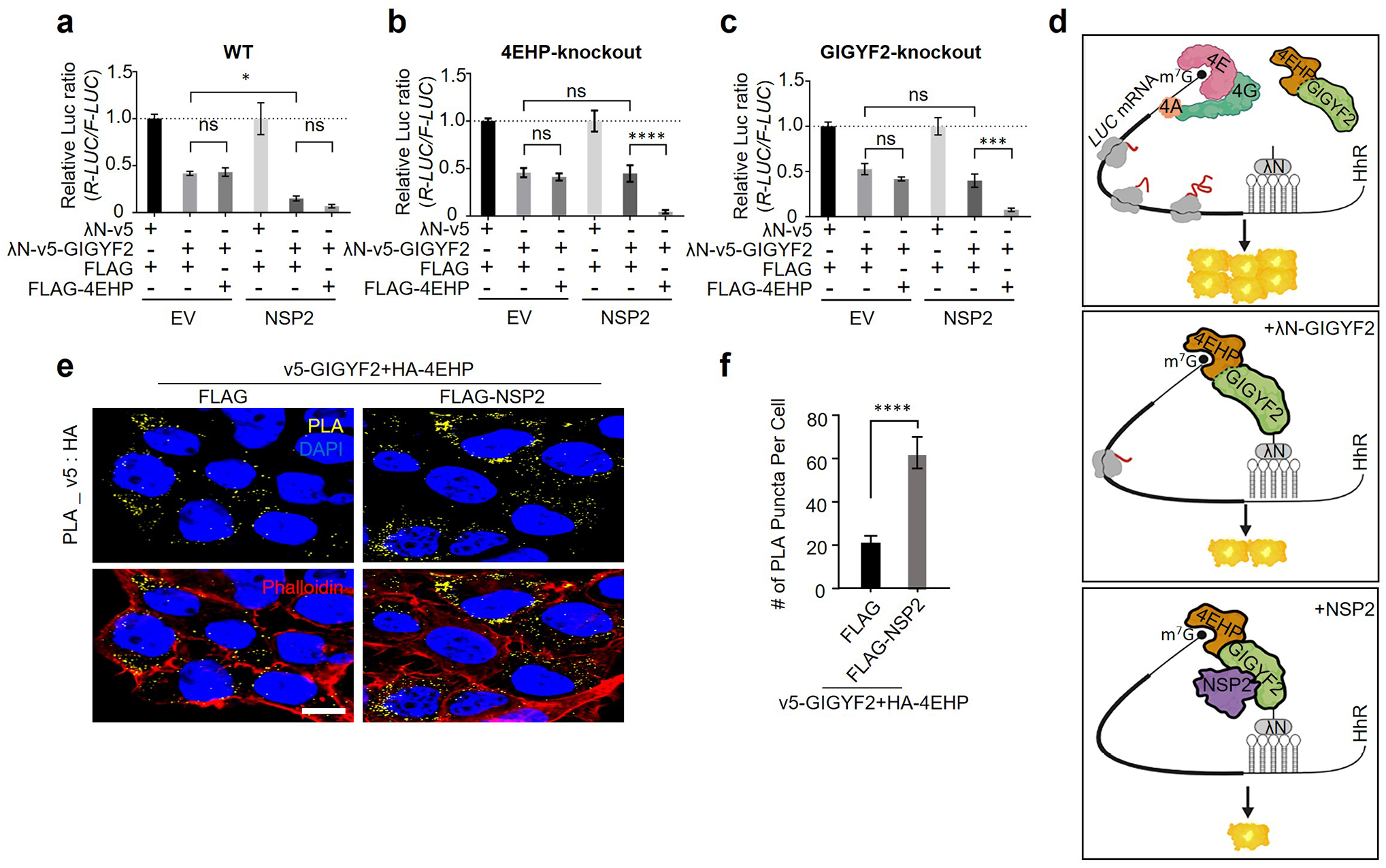
NSP2 augments GIGYF2/4EHP complex-mediated translational repression by enhancing the GIGYF2 interaction with 4EHP. **(a)** WT cells were co-transfected with vectors expressing either λN-v5-GIGYF2 or λN-v5 control, along with *R-Luc*-5BoxB-A114-N40-HhR and *F-Luc* (as control), in combination with FLAG-4EHP or FLAG-Empty plasmids. Dual-luciferase assay was performed 24 h post-transection. Data are presented as mean ± SD (n=3). *p<0.05, one-way ANOVA with Bonferroni’s post hoc test. (**b-c**) GIGYF2-tethering assay carried out in 4EHP-KO cells in (**b**) and GIGYF2-KO cells in (**c**). Data are presented as mean ± SD (n=3). ns= non-significant, **p<0.01, ***p<0.001, one-way ANOVA with Bonferroni’s post hoc test. (**d**) Graphic illustration of the GIGYF2/4EHP-mediated induction of translational repression by NSP2. (**e**) PLA assay for detection of GIGYF2-4EHP interactions in HEK293T cells transfected with vectors expressing v5-GIGYF2 and HA-4EHP together with FLAG-Empty or FLAG-NSP2. Cells were fixed and subjected to PLA using v5 and HA antibodies 24 h post-transfection. Scale bar= 10 µm. (**f**) Quantification of positive PLA signals from (e). The number of PLA signals from at least 20 cells was counted in each sample. n=5 independent experiments. Data are presented as mean ± SD (n=5). ****P< 0.0001, one-way ANOVA with Bonferroni’s post hoc test. See also Extended Data Fig. 2 and Fig. 3.

### The GIGYF2/4EHP complex represses IFN-β production

We previously reported that 4EHP suppresses *Ifnb1* mRNA translation^26^. 4EHP interacts with GIGYF2 and mediates GIGYF2-induced translational repression^23,33^. Thus, we examined the role of GIGYF2 in the regulation of IFN-β production. Toll-Like Receptor 3 (TLR3), was transiently expressed in WT, 4EHP-KO, and GIGYF2-KO HEK293 cells, which were then treated with poly(I:C), an angonist of TLR3 that stimulates IFN-β production. While 4EHP-KO cells produced ∼2.5-fold more IFN-β than WT cells, as expected^26^, a significantly more robust (∼5.5-fold) increase in IFN-β production was observed in GIGYF2-KO, compared with WT cells (Fig. 3a). Furthermore, consistent with the elevated IFN-β production, poly(I:C)-induced STAT1 phosphorylation was enhanced (∼3-fold) in 4EHP-KO HEK293 cells, an effect which was further augmented (∼6.5-fold) in GIGYF2-KO cells (Fig. 3b).

**Figure 3.**
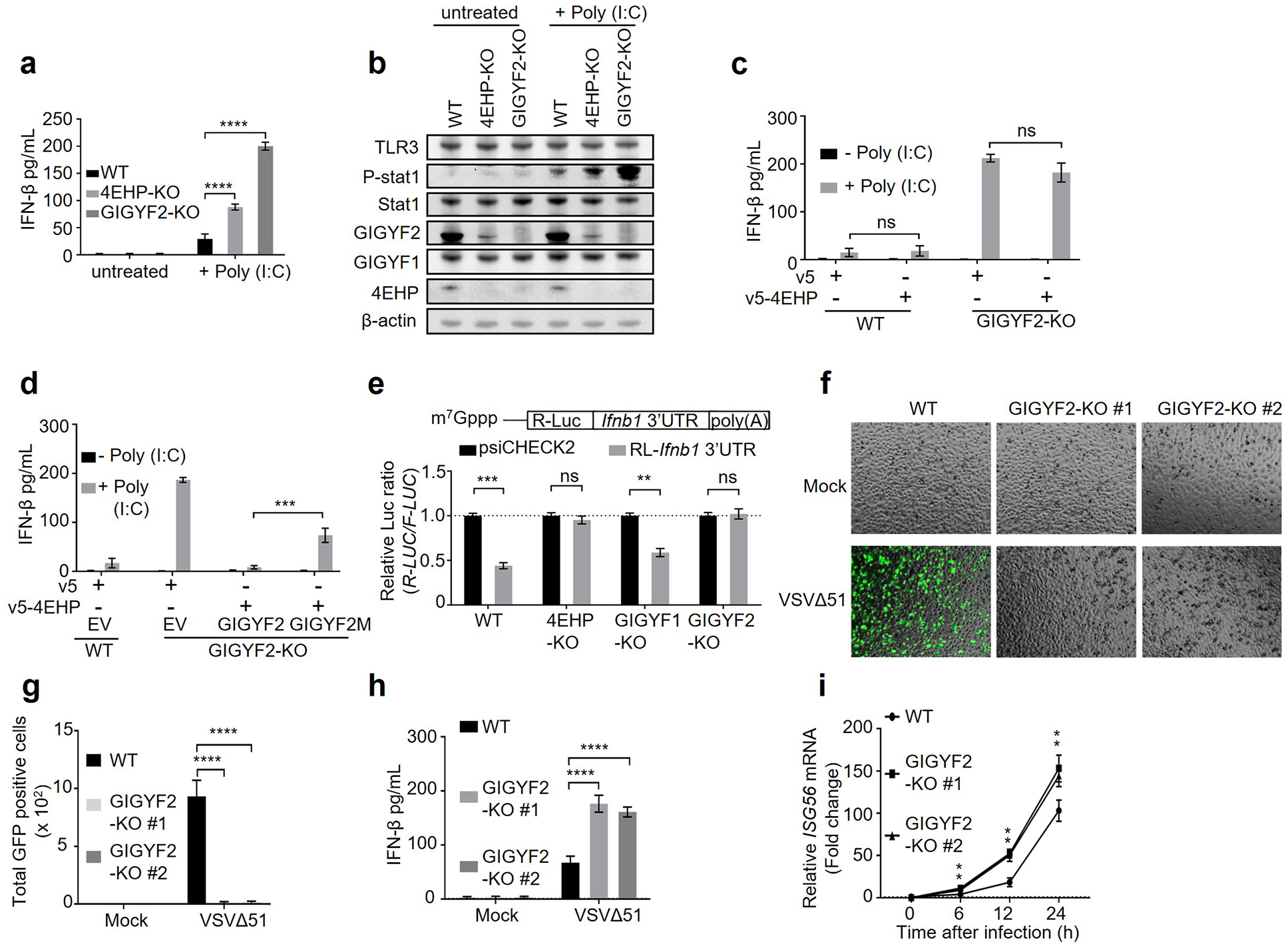
GIGYF2/4EHP complex formation is critical for repression of *Ifnb1* mRNA translation and enabling viral replication. **(a)** ELISA measurement of IFN-β production in WT, 4EHP-KO, and GIGYF2-KO HEK293 cells transiently expressing TLR3 following 6 h of treatment with poly(I:C). Data are presented as mean ± SD (n=3). ****P< 0.0001, one-way ANOVA with Bonferroni’s post hoc test. **(b)** Western blot analysis of cell lysates from (a). **(c)** WT and GIGYF2-KO HEK293 cells were transfected with the plasmids expressing v5-Empty or v5-4EHP. ELISA measurement of IFN-β production was performed following 6 h of poly(I:C) treatment. Data are presented as mean ± SD (n=3). ns=non-significant, two-tailed Student’s t-test. **(d)** v5-Empty or v5-4EHP expression plasmids were co-transfected with empty vector (EV) or plasmids expressing WT GIGYF2 or 4EHP-binding mutant GIGYF2 (Y41A, Y43A, M48A, L49A; GIGYF2M). IFN-β ELISA assay was performed following 6h of poly(I:C) stimulation. Data are presented as mean ± SD (n=3). ***P< 0.001, one-way ANOVA with Bonferroni’s post hoc test. **(e)** WT, 4EHP-KO, GIGYF1-KO, and GIGYF2-KO HEK293 cells were transfected with psiCHECK2-*R-Luc*-*Ifnb1* 3′ UTR reporter or the psiCHECK2 reporter (as control). *R-Luc* and *F-Luc* activities were measured 24 h after transfection. The *R-Luc*/*F-Luc* ratio in psiCHECK2-*R-Luc*-*Ifnb1* 3′ UTR reporter expressing cells was normalized to the psiCHECK2-expressing cells. Data are presented as mean ± SD (n=3). ns= non-significant, **P< 0.01, ***P< 0.001, two-way ANOVA with Bonferroni’s post hoc test. **(f-i)** WT or GIGYF2-KO#1-2 (2 independent sgRNAs) A549 cells were infected with mock or VSVΔ51-GFP (0.1 MOI). 12 h post-infection (h.p.i.), cells were subjected to: **(f)** visualization of VSVΔ51-GFP infection by fluorescence microscopy, **(g)** Quantification of the GFP positive cell number in (f) by ImageJ, **(h)** ELISA measurement of IFN-β production in the supernatant. **(i)** At the indicated time points post virus infection, *ISG56* mRNA levels were measured by RT-qPCR. *GAPDH* mRNA was used for normalization. Data are presented as mean ± SD (n=5). **P< 0.01, ***P< 0.001, ****P< 0.0001, one-way ANOVA with Bonferroni’s post hoc test. See also Extended Data Fig. 4 and Fig. 5.

Next, we tested whether GIGYF2 also suppresses IFN-β production in two lung epithelial cell lines, Calu-3 and A549, which are widely used in SARS-CoV-2 studies and respond to poly(I:C) stimulation with robust IFN-β production^34-37^. Upon poly(I:C) treatment of Calu-3 and A549 cells, IFN-β expression and STAT1 phosphorylation significantly increased in the 4EHP-depleted cells (2.5∼fold in Calu-3 and 1.8∼fold in A549 cells) and even more in GIGYF2-depleted cells compared to WT cells (6∼fold in Calu-3 and 6.1∼fold in A549 cells; Extended Data Fig. 4a-d). These data demonstrate that 4EHP and GIGYF2 repress IFN-β production and that GIGYF2 is a more potent repressor than 4EHP. Importantly, *Ifnb1* mRNA levels did not change in 4EHP- or GIGYF2-depleted cells compared to their control counterparts (Extended Data Fig. 4e-g).

Next, we investigated whether formation of the GIGYF2/4EHP complex is required for repression of IFN-β production. Rescuing expression of 4EHP, which is destabilized in GIGYF2-depleted cells^20^, failed to restore repression of IFN-β production in GIGYF2-KO HEK293 cells (Fig. 3c, Extended Data Fig. 5a). These data indicate that repression of IFN-β production by 4EHP requires the presence of GIGYF2. To determine whether direct interaction of 4EHP with GIGYF2 is required for repression of IFN-β production, we overexpressed 4EHP and WT GIGYF2 or a mutant version of GIGYF2 that does not bind to 4EHP (GIGYF2M)^20^, in GIGYF2-KO cells. Co-expression of 4EHP and WT GIGYF2 rescued the repression of IFN-β in GIGYF2-KO cells (Fig. 3d, Extended Data Fig. 5b). In contrast, co-transfection of 4EHP and GIGYF2M only partially (∼50%) rescued the IFN-β repression, indicating that formation of GIGYF2/4EHP complex is pivotal for efficient repression of IFN-β production.

The 3′ UTR of the *Ifnb1* mRNA plays a key role in 4EHP-mediated translational repression^26^. To investigate whether the *Ifnb1* mRNA 3′ UTR exerts its silencing effect via GIGYF2, we transfected a luciferase reporter (*R-Luc*) fused to the *Ifnb1* 3′ UTR into WT, 4EHP-KO, GIGYF1-KO, or GIGYF2-KO cells. Luciferase activity was repressed by 2-fold in WT and GIGYF1-KO cells, but not in 4EHP-KO and GIGYF2-KO cells (Fig. 3e), with no change in the abundance of *R-Luc* mRNA (Extended Data Fig. 5c). These data demonstrate that GIGYF2 and 4EHP mediate the translational silencing induced by *Ifnb1* mRNA 3′ UTR.

### GIGYF2 represses RNA virus replication

To assess a broader role of GIGYF2 in the antiviral immune response to RNA viruses through repression of IFN-β production, we used a GFP-tagged mutant variant (VSVΔ51) of the vesicular stomatitis virus (VSV). The deletion of methionine-51 (M51) in the matrix protein, renders the virus more sensitive to the IFN-mediated antiviral response^38^. We previously reported that 4EHP depletion inhibits the replication of VSVΔ51 by enhancing the production of IFN-β^26^. GIGYF2-KO significantly limited replication of GFP-tagged VSVΔ51 in A549 lung cells 12 hours post-infection(Fig. 3f & g). Following virus infection, expression of IFN-β (measured by ELISA) and the mRNA level of the IFN-stimulated gene 56 (ISG56, measured by RT-qPCR) were increased ∼2-fold as compared to WT cells (Fig. 3h & i) without a detectable change in *Ifnb1* mRNA levels (Extended Data Fig. 5d). These data support the conclusion that GIGYF2-depletion protects A549 cells from VSVΔ51-GFP infection, owing to robust IFN-β production and activation of IFN-induced antiviral pathways.

We next asked whether GIGYF2 also directly targets virus-induced activation of signaling pathways upstream of IFN-β. We first examined the impact of GIGYF2 or 4EHP depletion on virus RNA sensor-initiated signaling. We co-transfected GIGYF2-KO cells with a *F-Luc* reporter under the control of the minimum IFN-β promoter and constructs expressing constitutively active forms of key factors involved in RNA virus-induced signaling to mimic TLR3- and Retinoic acid inducible gene I (RIG-I)-like receptors (RLRs)-mediated signaling. GIGYF2-depletion did not affect IFN-β promoter activity mediated by upstream signaling pathways (Extended Data Fig. 5e). We also examined whether GIGYF2-depletion affects JAK-STAT, a key downstream signaling pathway activated by IFN-β. We transfected *F-Luc* reporter under the control of an IFN-sensitive response element (ISRE) promoter into WT, 4EHP-KO, or GIGYF2-KO HEK293 cells, followed by treatment with increasing doses of recombinant IFN-β protein. ISRE reporter activity was not affected by the removal of GIGYF2 or 4EHP-KO (Extended Data Fig. 5f). Neither did the removal of 4EHP or GIGYF2 affect recombinant IFN-β-induced STAT1 phosphorylation or ISG56 expression compared with WT cells (Extended Data Fig. 5g). These results demonstrate that the GIGYF2 and 4EHP-mediated antiviral immune response is a consequence of direct repression of *Ifnb1* mRNA translation and not via directly affecting the RNA virus sensors or signaling pathways downstream of IFN-β.

### SARS-CoV-2 NSP2 co-opts the GIGYF2/4EHP complex to repress *Ifnb1* mRNA translation

To investigate the potential role of NSP2 in the control of IFN-β expression, SARS-CoV-2 NSP2, NSP1, or Envelope (E) protein were expressed in HEK293 cells co-transfected with the *R-Luc* reporter construct fused to the *Ifnb1* 3′ UTR. The reporter expression was repressed ∼2-fold upon ectopic expression of NSP2, but not NSP1 or E protein (Fig. 4a, Extended Data Fig. 6a). To examine whether NSP2-mediated repression of *R-Luc*-*Ifnb1* 3′ UTR reporter requires the GIGYF2/4EHP complex, reporter activity was measured in WT, 4EHP-KO, and GIGYF2-KO cells expressing either GFP or NSP2 (Extended Data Fig. 6b). *R-Luc*-*Ifnb1* 3′ UTR was consistently repressed by 2-fold in the NSP2-transfected WT cells compared to the GFP-transfected WT cells (Fig. 4b), without affecting *R-Luc* mRNA levels (Extended Data Fig. 6c). However, reporter *R-Luc*-*Ifnb1* 3′ UTR silencing was relieved in 4EHP-KO and GIGYF2-KO cells, regardless of the expression of NSP2 (Fig. 4b). Thus, the NSP2-induced *Ifnb1* 3′ UTR-dependent repression requires the presence of 4EHP and GIGYF2. Notably, transient expression of NSP2 in WT HEK293 cells elicited a ∼40% and ∼48% increase in endogenous GIGYF2 and 4EHP protein levels, respectively (Extended Data Fig. 6b &d-e), without affecting the GIGYF2 and 4EHP mRNA levels (Extended Data Fig. 6f & g). This is likely due to the NSP2-induced enhanced GIGYF2 and 4EHP interaction (Fig. 2), which engenders mutual stabilization of both proteins^20^.

**Figure 4.**
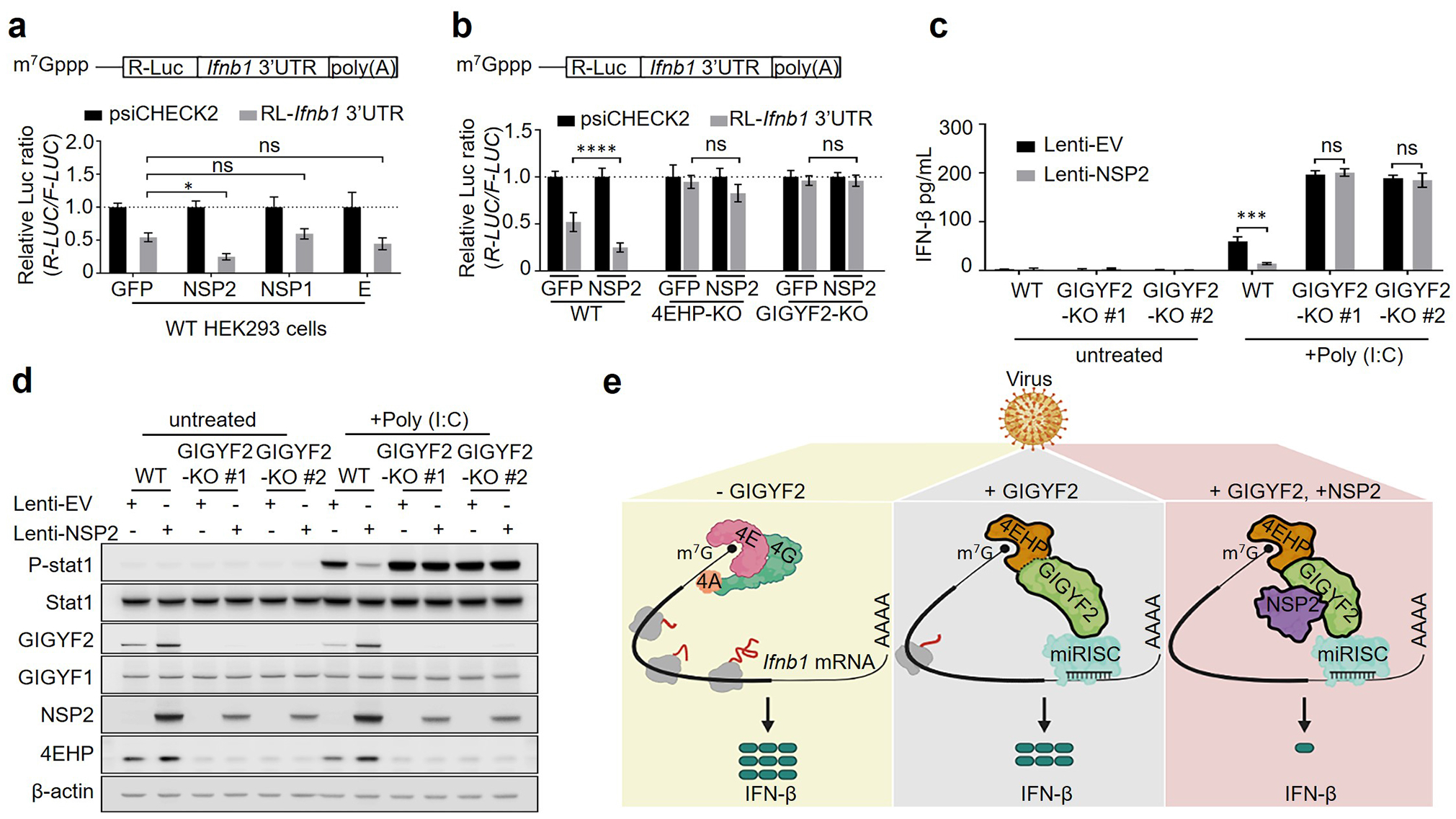
NSP2 augments the GIGYF2/4EHP complex-mediated translational silencing of *Ifnb1* mRNA. **(a)** 24 h post-transfection with GFP, NSP2, NSP1, or E protein, HEK293 cells were transfected with psiCHECK2-*R-Luc-Ifnb1* 3′ UTR reporter or the psiCHECK2 reporter (control). *R-Luc* and *F-Luc* activities were measured 24 h after the 2^nd^ transfection. The *R-Luc/F-Luc* ratio of psiCHECK2-*R-Luc-Ifnb1* 3′ UTR cells was normalized to the value for the psiCHECK2 cells as a percentage. Data are presented as mean ± SD (n=3). ns= non-significant, *p< 0.05, two-way ANOVA with Bonferroni’s post hoc test. **(b)** WT, 4EHP-KO, and GIGYF2-KO HEK293 cells were transfected with GFP or NSP2 expression plasmid for 24 h, followed by the 2^nd^ transfection with psiCHECK2-*R-Luc*-*Ifnb1* 3′ UTR reporter or the psiCHECK2 control reporter. 24h post-transfection the *R-Luc*/*F-Luc* ratio was measured as described in (a). **(c)** ELISA measurement of IFN-β and **(d)** western blot analysis of poly(I:C) treated WT and GIGYF2-KO A549 cells stably expressing either empty vector (EV) or NSP2 using lentiviral vector at 8 h post transfection. Data are presented as mean ± SD (n=3). ns= non-significant, ***p< 0.001, two-way ANOVA with Bonferroni’s post hoc test was used. **(e)** Graphic illustration of recruitment of the GIGYF2/4EHP repressor complex by NSP2 to silence IFN-β production in response to SARS-CoV-2 infection. GIGYF2/4EHP complex enables the miRISC-induced repression of the cap-dependent mRNA translation. Binding of NSP2 to GIGYF2 enhances the interaction of GIGYF2 with 4EHP, resulting in co-stabilisation of GIGYF2 and 4EHP and augmented translational repression of *Ifnb1* mRNA. See also Extended Data Fig. 6 and Fig. 7.

Next, we wished to confirm the above results in the lung epithelial cell line A549. We used lentiviral vectors to stably express NSP2 or control empty vector (EV) in WT and GIGYF2-KO A549 cells. Compared to the empty vector, ectopic NSP2 expression reduced the poly(I:C)-induced IFN-β production (∼4-fold; Fig. 4c) and STAT1 phosphorylation (Fig. 4d) in WT cells, without a significant impact on *Ifnb1* mRNA levels (Extended Data Fig. 7a). In stark contrast, NSP2 failed to repress IFN-β production or STAT1 phosphorylation in GIGYF2-KO cells (Fig. 4c & d). Notably, similar to HEK293 cells (Extended Data Fig. 6b), stable expression of NSP2 in A549 cells resulted in a 2-fold and 0.8-fold increase in GIGYF2 and 4EHP protein levels, respectively, without changes in *GIGYF2* and *4EHP* mRNA levels (Extended Data Fig. 7b-e). However, stabilization of 4EHP by NSP2 was not observed in NSP2-expressing GIGYF2-KO cells, because NSP2 directly interacts with GIGYF2, but not 4EHP.

Taken together, our data offer a mechanistic model in which NSP2 directly interacts with host GIGYF2 protein and enhances the interaction of GIGYF2 with 4EHP, resulting in stabilization of both GIGYF2 and 4EHP proteins. We show that NSP2 co-opts the GIGYF2/4EHP translational repression complex to suppress IFN-β production. The NSP2-induced enhancement of translational repression of *Ifnb1* mRNA and IFN-β production compromises antiviral immunity and is predicted to facilitate SARS-CoV-2 replication and exacerbate viral pathogenesis (Model; Fig. 4e).

### Functional and structural characterization of the interaction between NSP2 and GIGYF2

Next, we sought to understand the structural basis for the interaction between NSP2 and GIGYF2 using simulation programs, as the 3D structure of GIGYF2 is not resolved. Therefore, we queried human GIGYF2 (accession number: Q6Y7W6) against the Pfam database^39^ to predict its domains and folded regions. GIGYF2 is predicted to possess the known GYF motif (correctly designated between residues 557-601) in addition to a domain of unknown function spanning residues 744-940 (Fig. 5a). We performed secondary structure prediction on human GIGYF2 by the Garnier-Osgurthorpe-Robson IV (GOR4) method^40^. Consistent with the Pfam result, the region in GIGYF2 roughly spanning residues 750-950 is predicted with high confidence to fold into alpha helices (Extended Data Fig. 8a). To investigate the helical organization of GIGYF2 (744-940), we mapped this region back to a full-length three-dimensional model of human GIGYF2 as predicted by AlphaFold 2^16,41^. GIGYF2 (744-940) corresponds to a region in the model that is confidently predicted (Extended Data Fig. 8b & c) to fold into one singular, long alpha helix (Fig. 5a). Thus, our structural analysis of GIGYF2 reveals that this previously uncharacterized GIGYF2 region (744-940) corresponds to a singular alpha helix [hereafter referred to as the GIGYF2 long helix region (GIGYF2-LHR)]. We next sought to model the interaction between GIGYF2 and NSP2. A cryo-EM structure of SARS-CoV-2 NSP2 that modelled 68% of the total protein sequence^42^ was obtained from the Protein Data Bank (PDB code: 7MSX). There are natural variants of SARS-CoV-2 that possess glycine to valine point mutations at residues 262 and 265 in NSP2^43^. A recent study^42^ used an affinity purification mass-spectrometry (AP-MS) assay to investigate the changes in NSP2’s virus-host protein-protein interactome caused by naturally occurring mutations. Strikingly, an NSP2^G262V/G265V^ double mutant expressed in HEK293T cells failed to interact with the GIGYF2 and 4EHP^42^. A model for the interaction between GIGYF2 and NSP2 should therefore include and track the positions of G262/G265. To determine the 3D location of G262/G265 within folded NSP2, the intra-protein folding organization of NSP2 was predicted by analyzing its amino acid sequence with ColabFold^44^. NSP2 is predicted to contain five discrete folding units, four of which could be allocated to its cryo-EM structure (Fig. 5b, Extended Fig. 9a-d). G262/G265 occurred in a short alpha helix contained in a predicted folding unit spanning residues 251-375 (NSP2-R3) (Extended Data Fig. 9e). To model the interaction between GIGYF2 and NSP2 we performed protein-protein docking simulations between GIGYF2-LHR and either NSP2-R3 or the cryo-EM structure of NSP2 using HADDOCK 2.4^45^. Simulations between GIGYF2-LHR and NSP2-R3 yielded 8 docking models grouped into 2 clusters; cluster 2 had the lower and better HADDOCK score (Extended Data Fig. 10a). The best model of cluster 2 docked NSP2-R3 at the center of GIGYF2-LHR in a manner that positioned the G262/G265 containing helix proximal and parallel to GIGYF2-LHR (Extended Data Fig. 10c). Simulations between GIGYF2-LHR and the cryo-EM structure of NSP2 yielded 25 docking models grouped into 10 clusters, with cluster 6 performing with the lowest HADDOCK score (Extended Data Fig. 10b). The best model in this cluster also docked NSP2 at the center of GIGYF2-LHR with the helix containing G262/G265 directly adjacent to GIGYF2-LHR (Fig. 5c, Extended Data Fig. 10d). Overall, our docking simulations generated models for the interaction between GIGYF2 and NSP2 that were consistent with our structural analyses of these proteins and with the AP-MS data in the literature^42^.

**Figure 5.**
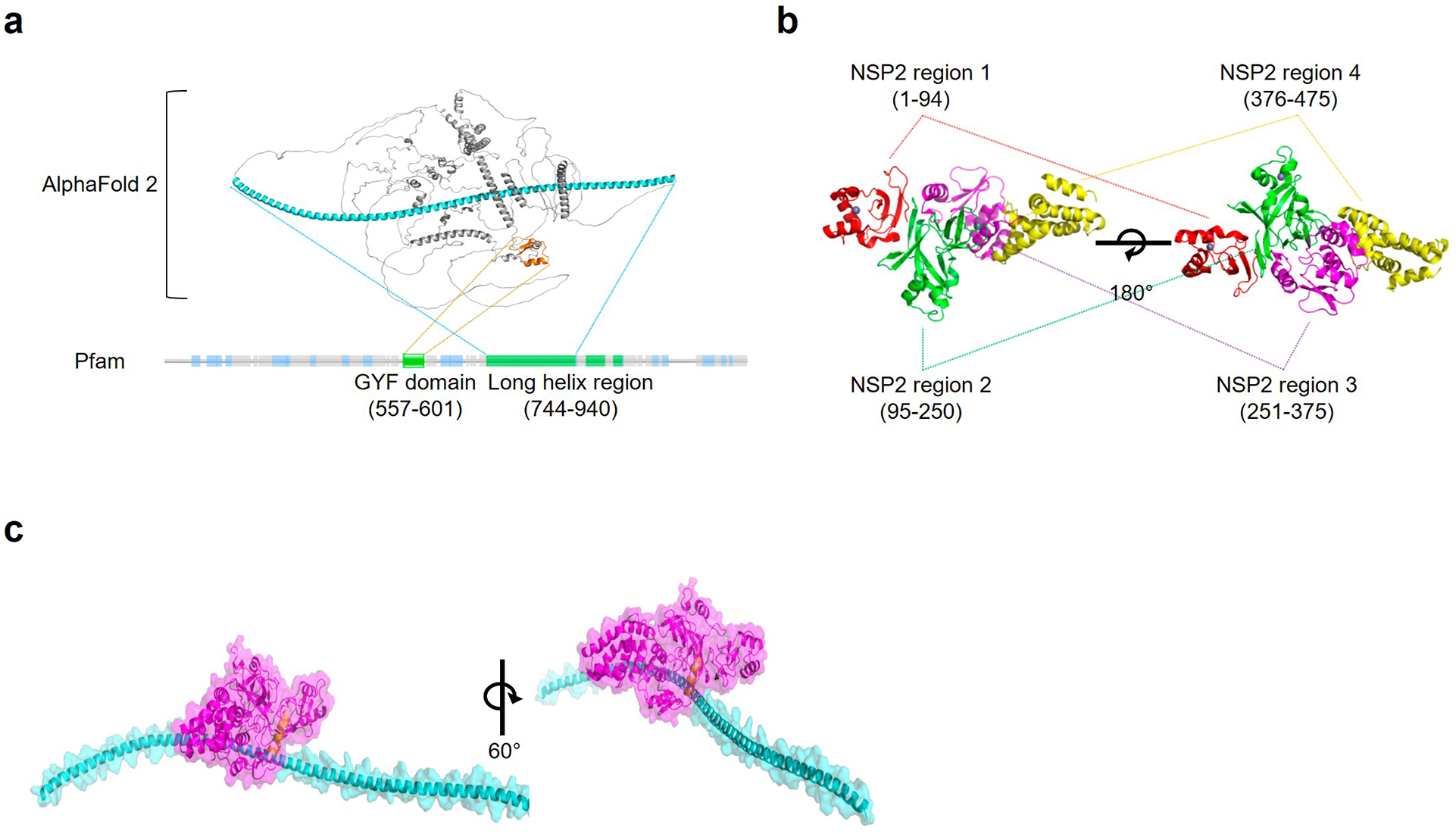
Functional and structural characterization of the interaction between NSP2 and GIGYF2. **(a)** Structural prediction and modeling of human GIGYF2 by Pfam and AlphaFold-2. The predicted three-dimensional structure of GIGYF2 was obtained from the AlphaFold Protein Structure Database and annotated in PyMOL: cyan – long helix region, orange – GYF domain, grey – rest of GIGYF2. The schematic of predicted domains and folding regions along the primary structure of GIGYF2 was generated by querying human GIGYF2 (accession number: Q6Y7W6) against the Pfam database. Grey – regions predicted to be disordered, blue – regions predicted to have low complexity, green – regions predicted to fold into known domains or secondary structures. **(b)** Cryo-EM structure of SARS-CoV-2 NSP2. The cryo-EM structure of NSP2 (4-104, 119-475, 491-505) was obtained from the Protein Data Bank (PDB code: 7MSX). Discrete folding regions within NSP2 were annotated in PyMOL: red – NSP2-R1, green – NSP2-R2, magenta – NSP2-R3, yellow – NSP2-R4. **(c)** Model of the interaction between GIGYF2-LHR and NSP2 by HADDOCK 2.4. The predicted model was annotated in PyMOL as follows: cyan – GIGYF2-LHR ribbon model superimposed over a GIGYF2-LHR 60% transparent surface model, magenta – NSP2 cryo-EM ribbon model superimposed over an NSP2 cryo-EM 60% transparent surface model, yellow – G262/G265 containing alpha helix in NSP2. See also Extended Data Fig. 8-10.

## Discussion

While a robust type I IFN-mediated antiviral innate immune response is indispensable for combating infections, an exacerbated response can result in pathological inflammation and tissue damage^46-49^. mRNA translational control mechanisms play a crucial role in maintaining the appropriate magnitude and duration of the immune response^49^. Our data show that the GIGYF2/4EHP complex inhibits translation of *Ifnb1* mRNA. We demonstrate that SARS-CoV-2 co-opt this mechanism through NSP2 protein, which is highly conserved among coronaviruses^42^ to impede upon the antiviral innate immune response. Thus, this mechanism is presumably utilized by other coronaviruses. Indeed, a previous study detected the interaction of SARS-CoV-1 encoded NSP2 with GIGYF2 and 4EHP^50^, indicating a common mechanism of impeding the host innate immune response by coronaviruses. We further revealed that another RNA virus (VSV) also employs the GIGYF2/4EHP complex to repress IFN-β production (Fig. 3)^26^. Thus, it appears that various RNA viruses use distinct approaches that converge on GIGYF2/4EHP translational repression complex to block the activation of the antiviral innate immune response.

Other SARS-CoV-2 proteins including NSP1 and NSP14 also dysregulate the host mRNA translation machinery^13,16,28,50,51^. NSP1 blocks the ribosomal entry site for host mRNAs but allows SARS-CoV-2 mRNA translation^52,53^. While viral RNA is protected, host mRNA is subjected to degradation. Thus, NSP1 non-specifically inhibits translation of host mRNAs including *Ifnb1*^53^ and results in depletion of labile antiviral factors such as Tyk2 and STAT2^54^. NSP14 also inhibits global mRNA translation, which likewise involves the shutdown of ISGs expression^51^. In contrast, we showed that NSP2 associates with the GIGYF2/4EHP complex to repress translation of *Ifnb1* mRNA, but it is highly likely that this mechanism also affects the expression of other important cytokines that promote antiviral response.

The N-terminal region of GIGYF2 encodes several important protein binding motifs, including the 4EHP-binding motif^20^, DDX6-binding motif, and the GYF domain which interacts with the Pro-Pro-Gly-hydrophobic motif (PPGL)^20,55^. We described a candidate NSP2 binding domain at the LHR of GIGYF2. The 3D structure of the LHR and its interaction with NSP2 will be instrumental for a better understanding of the molecular basis of the proposed NSP2/GIGYF2/4EHP complex. Lastly, identifying the binding motif on NSP2 and GIGYF2 LHR could inform the development of recombinant peptides or small molecules to abolish the interaction of NSP2 with GIGYF2. It is noteworthy that a functional CRISPR library screen identified GIGYF2 and 4EHP as essential host proteins for efficient SARS-CoV-2 viral replication^56^. The knowledge of the mechanism of action of NSP2-mediated IFN suppression via the 4EHP/GIGYF2 complex is of considerable value for devising drugs to combat future infections of SARS-CoV-2, and of other known and unknown coronaviruses.

## METHODS

### Cell lines and culture conditions

HEK293T (Thermo Fisher Scientific, Waltham, MA) cells were cultured in DMEM (Dulbecco’s Modified Eagle Medium) supplemented with 10% fetal bovine serum (FBS) and 1% penicillin/streptomycin (P/S) (Wisent Technologies). A549 (ATCC), were cultured in RPMI, also supplemented with 10% FBS and 1% P/S. Calu-3 (ATCC) were cultured in EMEM (Eagle Minimal Essential Medium) supplemented with 20% FBS and 1% P/S. WT, 4EHP-knockout and GIGYF2-knockout HEK293 cells were maintained in DMEM supplemented with 10% FBS, 1% P/S, 100 µg/mL zeocin (Thermo Fisher Scientific, R25001), and 15 µg/mL blasticidin (Thermo Fisher Scientific, R210-01)^17^. All cells were cultured at 37°C, in a humidified atmosphere with 5% CO_2_.

### Antibodies, siRNAs, shRNAs, and plasmids

The following antibodies were used: rabbit anti-eIF4E2 (Genetex, GTX103977 and GTX64395), rabbit anti-GIGYF1 (Bethyl Laboratories, A304-132A), rabbit anti-GIGYF2 (Bethyl Laboratories, A303-732A), sheep anti-SARS-CoV-2-NSP2 (MRC PPU reagents and Services, DA105), mouse anti-β-actin (Sigma, A5441), rabbit anti-STAT1 (Cell Signaling, 14994), rabbit anti-phospho-STAT1 (Tyr701; Cell signaling, 7649), rabbit anti-TLR3 (Cell signaling, 6961), rabbit anti-v5 (abcam, ab9116), mouse anti-FLAG (abcam, ab49763). The following shRNAs were used: Non-Targeting Control shRNA (Sigma, SHC002), EIF4E2 shRNA#1 (sh4EHP#1) (Sigma, TRCN0000152006), EIF4E2 shRNA#2 (sh4EHP#2) (Sigma, TRCN0000280916) and EIF4E2 shRNA#3 (sh4EHP#3) (Sigma,). GIGYF2 shRNA#1 (shGIGYF2#1) (Sigma, TRCN0000135151), GIGYF2 shRNA#2 (shGIGYF2#2) (Sigma, TRCN0000138937) and GIGYF2 shRNA#3 (shGIGYF2#3) (Sigma, TRCN0000135088).

Plasmids encoding *Firefly luciferase* (*F-Luc*) driven by either the ISRE promoter (ISRE-Luc) or the *Ifnb1* promoter (IFN-β–Luc) were used for the reporter assay. The pRL-TK vector (Promega, E2241) encoding *Renilla luciferase* (*R-Luc*) was used as a transfection control. TLR-3 expressing plasmid was used to transfect HEK293 cells transiently. The plasmids encoding human RIG-I, MAVS, TBK1, TRIF, and IRF3 have been previously described^57^. The pLenti-CMV-GFP-Puro (Addgene, 17448), pLenti-CMV-Luc-Puro (Addgene, 17477), Lenti-4EHP (Sigma, TRCN0000474313), and pLenti-X2-Zeo-DEST (Addgene, 21562) were used to generate cells which stably express GFP, luciferase,4EHP, and NSP2, respectively. The Lenti-Cas9-Blast (Addgene, plasmid 52962) and pLenti-CRISPRv2 (Addgene, plasmid 52961) were used to generate the knockout.

### Generation of knockout cell lines by CRISPR-Cas9

CRISPR-Cas9-mediated genome editing of Flp-In T-REx HEK293 cells was performed as previously described^58^. The oligodeoxynucleotides encoding sgRNAs for targeting the coding region of the gene of interest are listed in Extended Data Table. 1. Briefly, the forward and reverse strand oligodeoxynucleotides were annealed and ligated into pSpCas9(BB)-2A-GFP (Addgene, PX458, Plasmid #48138) linearized with BbsI (Thermo Fisher Scientific, ER1011). After transformation, the guide sequence containing pSpCas9(BB)2A-GFP plasmids were isolated and sequence-verified. To generate gene knockout Flp-In T-REx HEK293 cells, 130,000 cells were transfected with the corresponding guide sequence containing pSpCas9(BB)-2A-GFP plasmid. 24 h after transfection, GFP positive cells were-single cell sorted by FACS into two 96-well plates and cultivated until colonies were obtained.

The 4EHP-KO and GIGYF2-KO A549 cells were generated in two steps. Firstly, the A549 lenti-Cas9 cells were generated by infecting the parental cells with lenti-Cas9 lentiviral particles, followed by selection of the infected cells with 10 ug/mL blasticidin (Thermo Fisher Scientific, R210-01). The Cas9 stable A549 cells were then infected with packaged lentivirus pLenti-CRISPRv2 expressing small guide RNA (sgRNA) of interest, followed by treatment with 10 ug/mL puromycin (Bioshop, PUR333.500). The sequence of sgRNAs for targeting the coding region of the gene of interest are listed in Extended Data Table 1.

Clonal cell lines were analyzed by WB for the absence of the protein and further analyzed for indel mutations within the targeted alleles by PCR. PCR products were cloned using the Zero Blunt PCR Cloning Kit (Thermo Fisher Scientific, K270040) and 10 clones were sequenced per cell line. The primers used for the PCR genotyping are listed in Extended Data Table 1.

### Lentivirus production

Lentivirus pseudovirions were produced by transfecting HEK293T cells using Lipofectamine 2000 and 10 μg shRNAs, lenti-Cas9-Blast, pLenti-CRISPRv2 inserted with CRISPR guide RNA (gRNA) for gene knockout plasmids, or Lenti-ORF plasmid, 6 μg psPAX2 (Addgene, plasmid 12260) and 4 μg pMD2.G (Addgene, plasmid 12259). 48 h post-transfection, cell culture supernatant was collected from which pseudovirions were purified by ultracentrifugation (32,000 rpm) for 2 h. The pelleted virus was resuspended into DMEM medium. Virus titer was adjusted to 5 multiplicity of infection (MOI).

### VSVΔ51-GFP virus infections

GFP-expressing VSVΔ51-GFP was previously described^59^. Virus titer was determined using a standard plaque assay protocol^60^. Virus replication was assessed *via* fluorescence microscopy. Viral MOIs used in each assay is described in the corresponding figure legends.

### Pattern Recognition Receptors (PRRs) ligands treatment and ELISA

Calu-3, HEK293 and A549 cells were seeded at 50-60% confluency. Cells were treated for 6 h with the appropriate concentration of either poly(I:C) (Sigma, P1530) or High Molecular Weight (HMW) poly(I:C) (InvivoGen, tlrl-pic) using Lipofectamine2000. IFN-β amounts in the culture supernatant were measured by human IFN-β ELISA kit (R&D Systems, DIFNB0) according to the manufacturer’s protocol.

### RNA extraction and RT-PCR

Cells were harvested and RNA was isolated using mammalian total RNA isolation kit (Sigma, RTN70-1KT). Following the Superscript III reverse transcriptase protocol (Invitrogen), equal amounts of total RNA (1 μg) were used for reverse transcription with 100 ng random primers. mRNA abundance was estimated *via* real-time PCR system (Mastercycler Realplex, Eppendorf) using SYBR Green master mix (Bio-rad). All primers are listed in Extended Data Table. 1.

### Plasmid construction

Synthesized viral coding sequences were incorporated into Gateway-compatible Entry vectors; pDONR207 SARS-CoV-2 NSP1 (Addgene, 141255), pDONR223 SARS-CoV-2 NSP2 (Addgene, 141256) and expression clones with N-terminal fusion tags were produced simply by Gateway cloning (Gateway™ LR Clonase™ II Enzyme mix, Invitrogen, 11791020). The Lenti-NSP2 was constructed by using the destination vector, pLenti-X2-Zeo-DEST (749-3) [a gift from Eric Campeau (Addgene, 21562)] and the donor vector, pDONR223 SARS-CoV-2 NSP2. The Lenti-NSP2 plasmid was used to generate NSP2 stable cell lines by following the lentivirus production method section.

### Dual Luciferase reporter assays

The *Ifnb1* promoter induced luciferase assay were descripted before^26^. Briefly, the IFN-β–Luc was co-transfected with pRL-TK in wild type and 4EHP-KO HEK293 cells using Lipofectamine 2000 according to the manufacturer′s protocol (Invitrogen). 24 h after transfection, cells were lysed. Lysates were used to measure the activity of *F-Luc* and *R-Luc via* the Dual-Luciferase Reporter Assay System (Promega, E1960) in a GloMax 20/20 luminometer (Promega, USA) according to the manufacturer’s instructions.

We adapted a previously used ISRE promoter-induced Luciferase assay in this study^26^. The ISRE-Luc was co-transfected with pRL-TK in control, 4EHP-KO and GIGYF2-KO HEK293 cell lines. 16 h post-transfection, cells were treated with different concentrations of recombinant human IFN-β (R&D Systems, 8499-IF-010) for 12 h. Relative *F-Luc* activity compared to *R-Luc* was quantified using a dual luciferase assay (Promega, E1960).

The psiCHECK-2 control vector (Promega, C8021) and the psiCHECK-RL-*Ifnb1* 3′ UTR vector were described before^26^. Briefly, the 3′ UTR sequence of human *Ifnb1* mRNA was inserted into the XhoI and NotI restriction sites in the psiCHECK*-*2 vector downstream of the *Renilla luciferase* ORF^26^. WT, 4EHP-KO or GIGYF2-KO HEK293 cells (150,000 cells/well) were co-transfected with either 10 ng psiCHECK*-*2 reporter (as control) or a construct of luciferase with full length *Ifnb1* 3′ UTR (psiCHECK2-RL-*Ifnb1* 3′ UTR) using Lipofectamine 2000. 24 h post-transfection, cells were lysed, followed by dual-luciferase assay. *R-Luc* values were normalized against *F-Luc* levels for each sample.

For the tethering assay, we used the λN-BoxB tethering approach^32^. Briefly, a *R-Luc* reporter containing five BoxB hairpins in its 3′ UTR was used. The 3′ end of the reporter contains a self-cleaving hammerhead ribozyme (HhR) to generate an internalized poly(a) stretch to prevent deadenylation and subsequent degradation^31^. The reporter was co-transfected along with the construct encoding a fusion of the protein of interest ORF, like GIGYF2, to a λN peptide, which allows GIGYF2 to bind to reporter BoxB elements. HEK293 cells were co-transfected with the constructs expressing either λN-V5 control or λN-V5 fused with protein of interest along with *R-Luc*-*5boxB*-A114-N40-HhR or *R-Luc*-A114-N40-HhR, and *F-Luc*, followed by the dual-luc assay after 24 h transfection. The silencing effect of *R-Luc* (normalized by *F-Luc*) mediated by the protein of interest was examined by comparing the dual-luciferase activities of the cells.

### Proximity Ligation Assay (PLA)

Proximity ligation assay (PLA) using Duolink reagents (Sigma, DUO92101) was performed according to the manufacturer’s recommendations. Briefly, cells were fixed with 4% PFA-sucrose for 15 min and permeabilized by PBS containing 0.1% Triton X-100 for 15 min. Cells were blocked in Duolink blocking solution for 1 h at 37°C and incubated with primary antibodies overnight at 4°C. Cells were washed by Wash Buffer A before incubation with PLA probe for 1 h at 37 °C, followed by ligation for 30 min at 37°C. PLA signals were amplified using Amplification Buffer for 100 min at 37°C, followed by washing with Wash Buffer B and mounting onto the slide glass before Airyscan microscopic imaging (Zeiss).

### Quantification and Statistical Analyses

Statistical tests were performed using Prism 6 (GraphPad). Error bars represent standard deviation (SD) from the mean. Number of independent replicates (≥3) and the statistical analysis used for each assay is described in the relevant figure legends. P values <0.05 were considered significant.

## Acknowledgments

We thank Samer Girgis and Trista Lou for technical assistance. This work was supported by grants from the Canadian Institutes of Health Research (CIHR; FDN-148423) to N.S. and the Terry Fox Research Institute (1073) to N.S. and T.A., and the Biotechnology and Biological Sciences Research Council (BB/W008165/1) to S.M.J. X.Z. is supported by funding from the China Scholarship Council (CSC). J.H.C is supported by the Basic Science Research Program through the National Research Foundation of Korea funded by the Ministry of Education (2020R1A6A3A03040141). S.M.J. is supported by the Patrick Johnston Research Fellowship, Queen’s University, Belfast. The graphical abstract was generated with BioRender.com.

## AUTHOR CONTRIBUTIONS

X.Z., J.H.C., and C.Z. performed the majority of the experiments. D.L.D. performed the *in silico* modeling. J.L., R.L., Q.L., P.N. contributed to the experiments. X.Z., N.S., and S.M.J., conceptualized the study. X.Z., N.S., J.H.C, and S.M.J. designed the experiments. N.S. and S.M.J. oversaw the study. N.S., S.M.J., J.H.C., X.Z., D.L.D., T.A., T.D., R.L., and S.W. wrote the manuscript. All authors reviewed and edited the manuscript.

