## Supplemental Figures and Table for "SARS-CoV-2 impairs interferon production via NSP2-induced repression of mRNA translation"

#### Extended Data Figure. 1

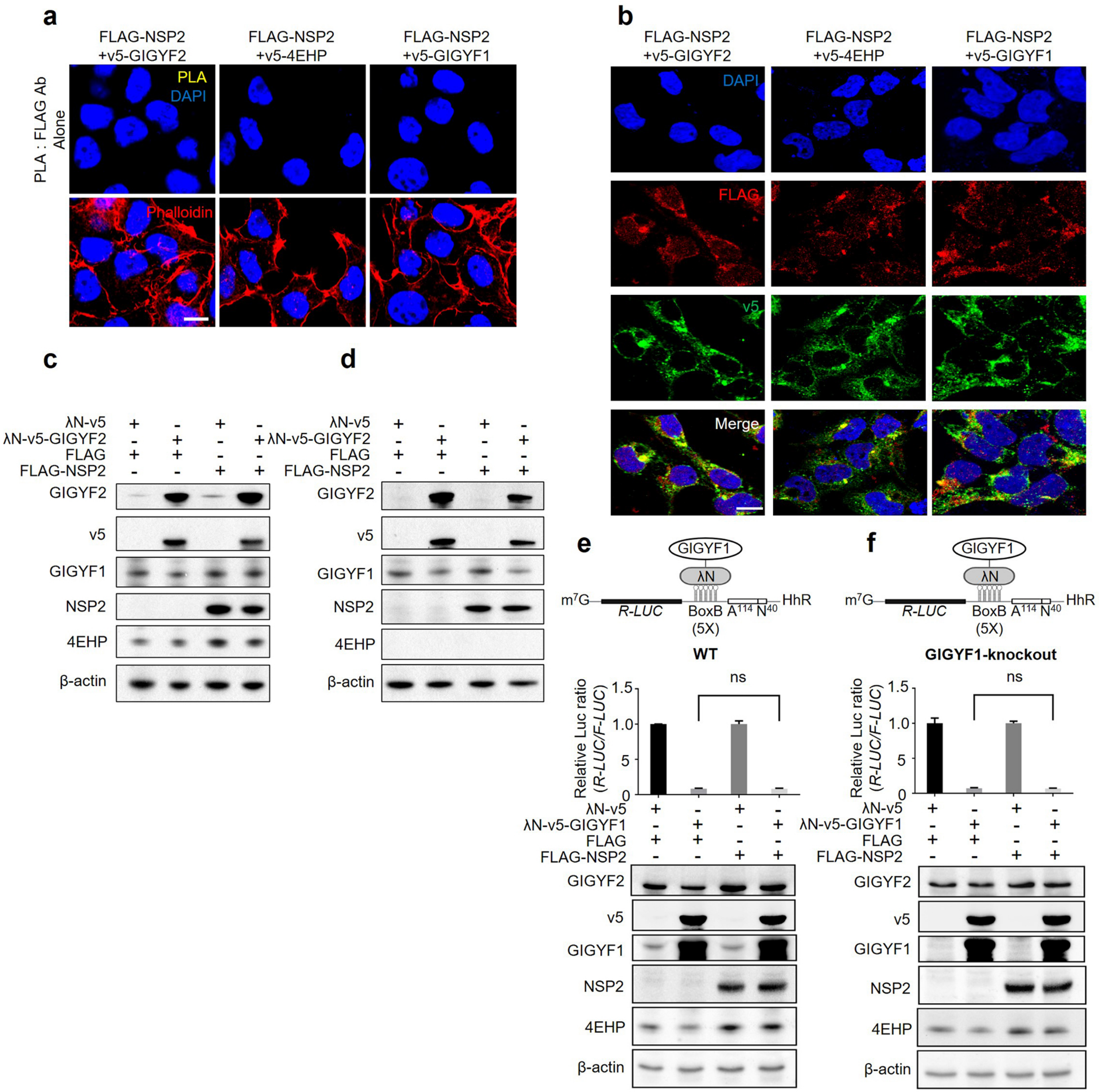

**Extended Data Figure 1. NSP2 does not enhance GIGYF1-mediated translation repression; related to Figure 1.** (**a**) Negative controls represent PLA performed with a single antibody (PLA: FLAG ab alone). Scale bar= 10 μm, n=5 independent replicates. (**b**) Immunofluorescence staining with the indicated antibodies in cells shown in Fig. 1a. Scale bar= 10 μm. (**c & d**) Western blotting with the indicated antibodies using lysates from the cell shown in Fig. 1c and Fig. 1d, respectively. (**e**) GIGYF1 tethering dual-luciferase assay in the presence or absence of FLAG-NSP2 was performed in WT HEK293 cells (upper panel) and cell lysate was used for western blotting (lower panel). Data are presented as mean ± SD (n=3). ns= non-significant, one-way ANOVA with Bonferroni's post hoc test. (**f**) GIGYF1 tethering dual-luciferase assay (upper panel) and western blot (lower panel) analysis were performed in GIGYF1-KO cells in the presence or absence of FLAG-NSP2. Data are presented as mean ± SD (n=3). ns= non-significant, one-way ANOVA with Bonferroni's post hoc test.

### Extended Data Figure. 2

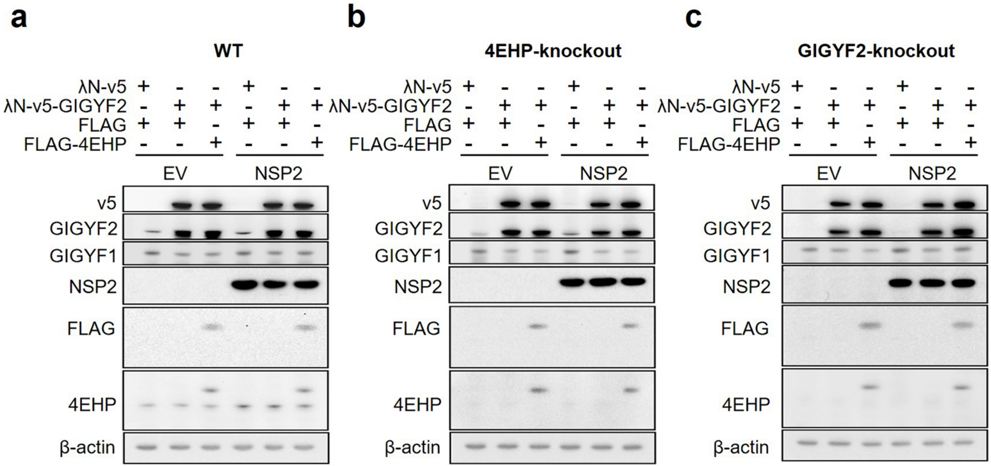

**Extended Data Figure 2. Western blotting analysis of the GIGYF2 tethering experiments, related to Figure 2.** (**a-c**) Western blotting with the indicated antibodies using lysates from the cells shown in Fig. 2a-c.

### Extended Data Figure. 3

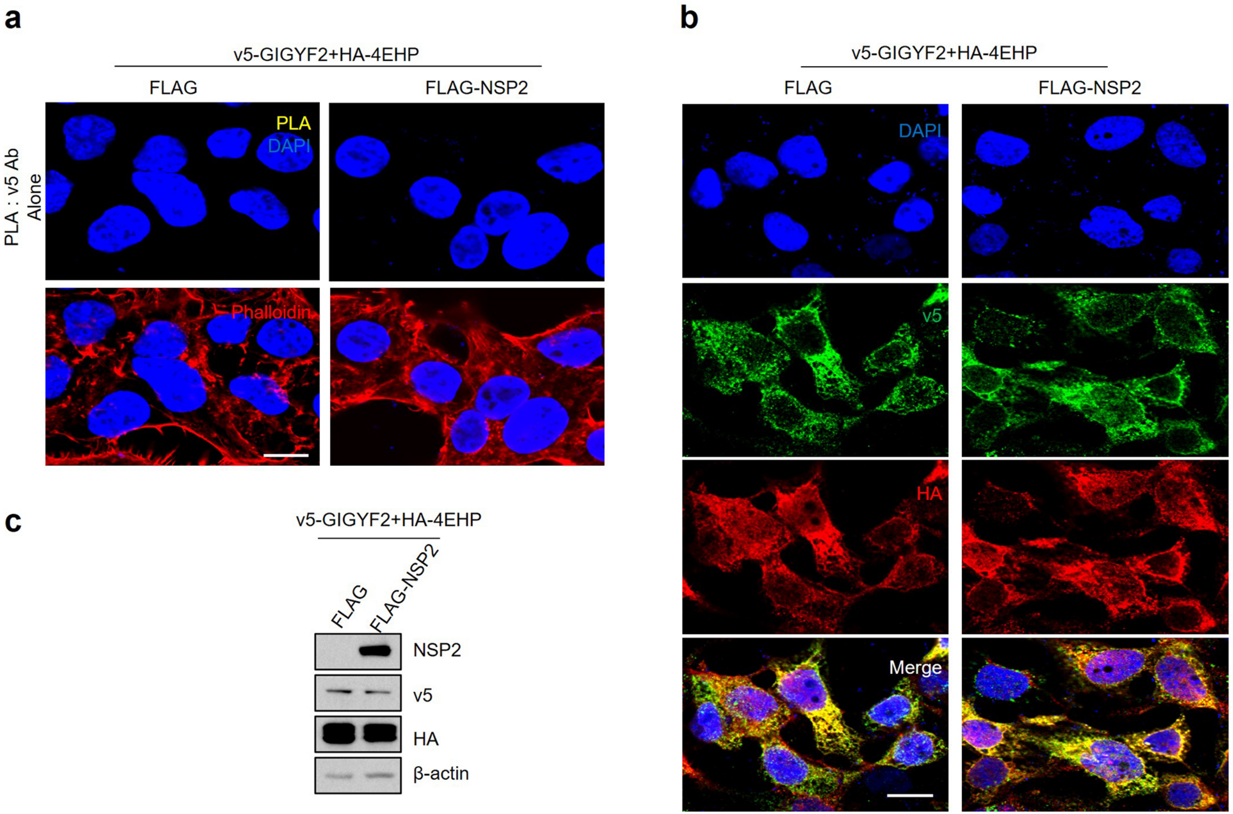

**Extended Data Figure 3. Immunofluorescence visualisation of markers used in PLA experiments; related to Figure 2.** (**a**) Negative controls represent PLA punctate performed with a single antibody (PLA: v5 ab alone). Scale bar= 10 μm, n=5 independent experiments. (**b**) Immunofluorescence staining performed with cells shown in Fig. 2e. Scale bar= 10 μm. (**c**) Western blot analysis of cells shown in Fig. 2e.

#### Extended Data Figure. 4

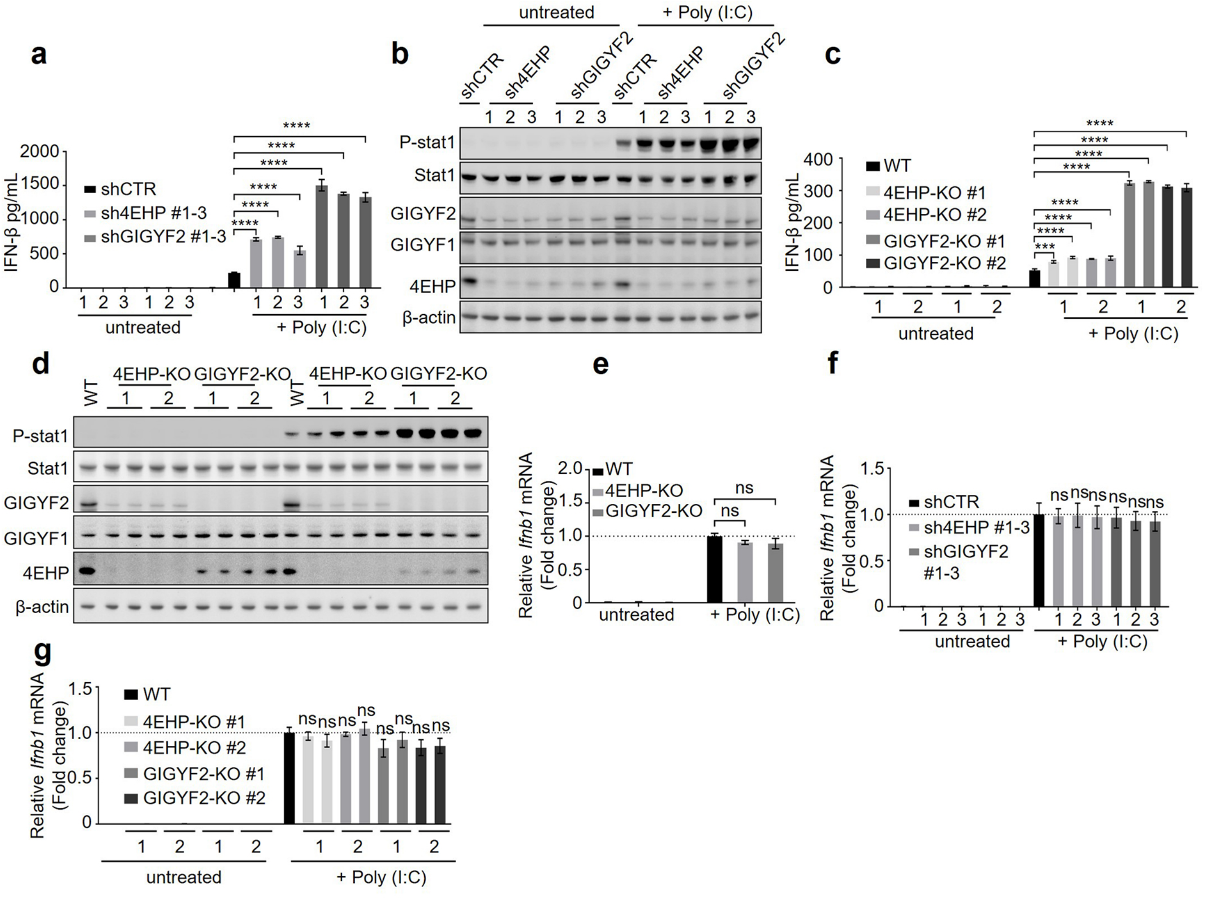

**Extended Data Figure 4. GIGYF2/4EHP complex represses IFN-β production without affecting *Ifnb1* mRNA level; Related to Figure 3. (a)** ELISA measurement of IFN-β production in shCTR, sh4EHP#1-3 and shGIGYF2#1-3 in Calu-3 cells 6 h post-treatment with poly(I:C). Data are presented as mean ± SD (n=3). ****$P < 0.0001$, one-way ANOVA with Bonferroni's post hoc test. **(b)** Western blot analysis of cell lysates from (a). **(c)** ELISA measurements of IFN-β production in WT, 4EHP-KO#1-2, and GIGYF2-KO#1-2 A549 cells after 6 h treatment with poly(I:C). Data are presented as mean ± SD (n=3). ***$P < 0.001$, ****$P < 0.0001$, one-way ANOVA with Bonferroni's post hoc test. **(d)** Western blot analysis of cell lysates from (c). **(e)** RT-qPCR analysis of WT, 4EHP-KO, and GIGYF2-KO HEK293 cell lines transiently expressing TLR3 (treatment described in Fig. 3a) 6 h post-treatment with poly(I:C). **(f)** RT-qPCR analysis of shCTR, sh4EHP#1-3, and shGIGYF2#1-3 Calu-3 cells treated with poly(I:C). **(g)** RT-qPCR analysis of WT, 4EHP-KO#1-2, and GIGYF2-KO#1-2 A549 cell lines 6 h post-treatment with poly(I:C).

### Extended Data Figure. 5

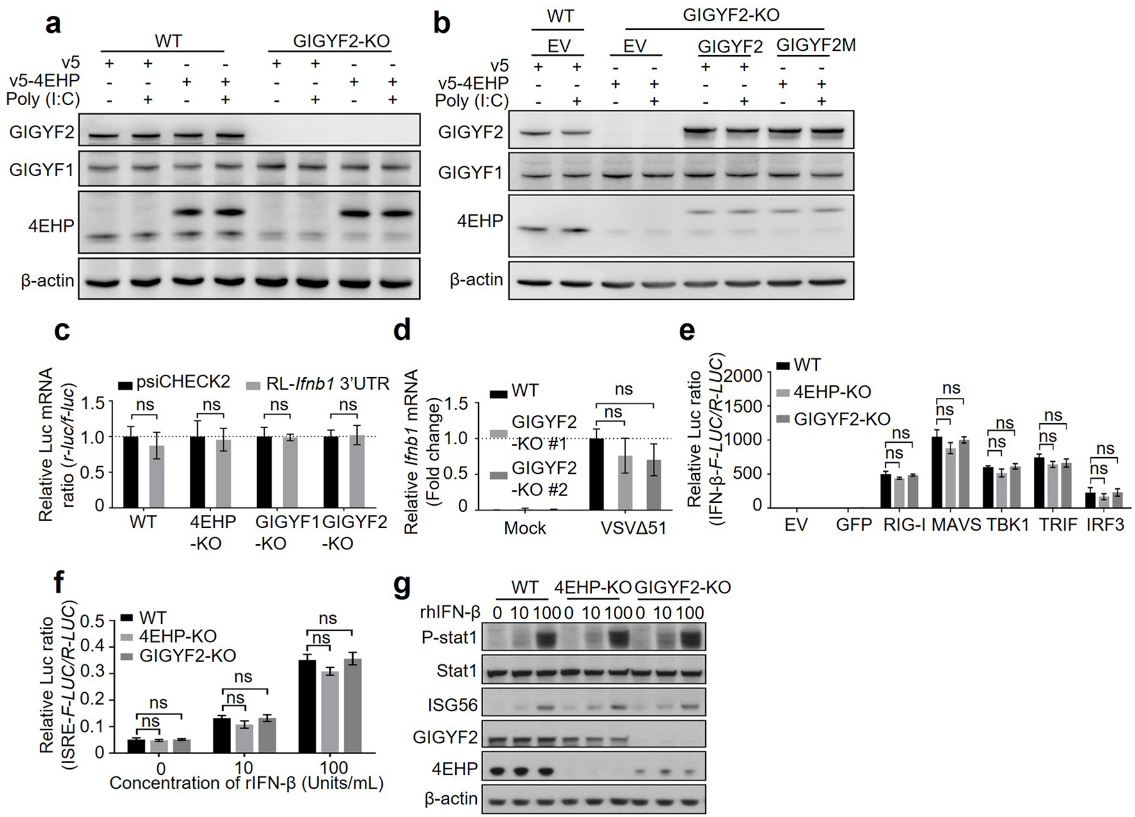

**Extended Data Figure 5. GIGYF2/4EHP complex does not directly affect the signalling pathways upstream and downstream of IFN-β; Related to Figure 3. (a)** Western blot analysis of lysates from Fig. 3c. **(b)** Western blot analysis of lysates from Fig. 3d. **(c)** RT-qPCR measurement of the *R-Luc* mRNA levels described in Fig. 3e. The *Luciferase* mRNA levels in the empty vector were used for normalization. Data are presented as mean ± SD (n=3). ns=non-significant, two-way ANOVA with Bonferroni's post hoc test. **(d)** RT-qPCR analysis of *Ifnb1* mRNA in the WT or GIGYF2-KO#1-2 A549 cells following VSVΔ51-GFP virus infection, as described in Fig. 3f. **(e)** WT, 4EHP-KO, or GIGYF2-KO HEK293 cells were co-transfected with *Ifnb1* promoter-driven *F-Luc* (*IFN-β-Luc*) and *R-Luc* expression plasmids 24 h post-transfection with empty vector, GFP, RIG-I, MAVS, TBK1, TRIF, or IRF3. Luciferase activities were measured 16 h after the 2$^{nd}$ transfection. **(f)** The WT, 4EHP-KO, and GIGYF2-KO HEK293 cells were co-transfected with both *ISRE-F-Luc* and *R-Luc* plasmids. 16 h post-transfection, cells were treated with indicated concentrations of recombinant human IFN-β for 12 h. The *ISRE-F-Luc*/*R-Luc* ratios were quantified by dual luciferase assay. ns=non-significant. **(g)** Western blot analysis of cell lysates from (f).

### Extended Data Figure. 6

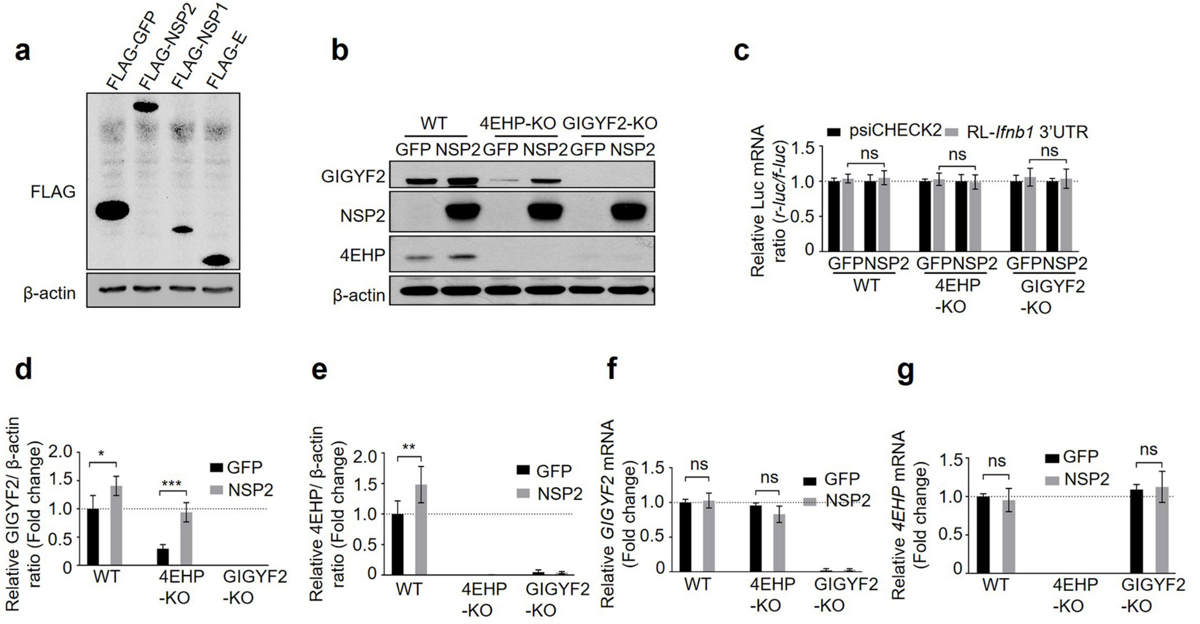

**Extended Data Figure 6. Ectopic expression of NSP2 enhances endogenous GIGYF2 and 4EHP protein expression without altering the *GIGYF2* and *4EHP* mRNA abundance; Related to Figure 4. (a)** Western blot analysis of cell lysates from Fig. 4a. (**b**) Western blot analysis of the indicated proteins in cell lysates from Fig. 4b. **(c)** RT-qPCR measurement of mRNA levels of the luciferase reporters described in Fig. 4b. The luciferase mRNA levels in empty vector were used for normalization. **(d-e)** Quantitation of expression of endogenous GIGYF2 (**d**) or 4EHP (**e**). β-actin expression was used as internal control. **(f-g)** RT-qPCR analysis of *GIGYF2* mRNA (**f**) or *4EHP* mRNA (**g**) levels. The WT HEK293 cells overexpressing GFP was used for normalization. Data are presented as mean ± SD (n=3). ns=non-significant, two-way ANOVA with Bonferroni's post hoc test.

#### Extended Data Figure. 7

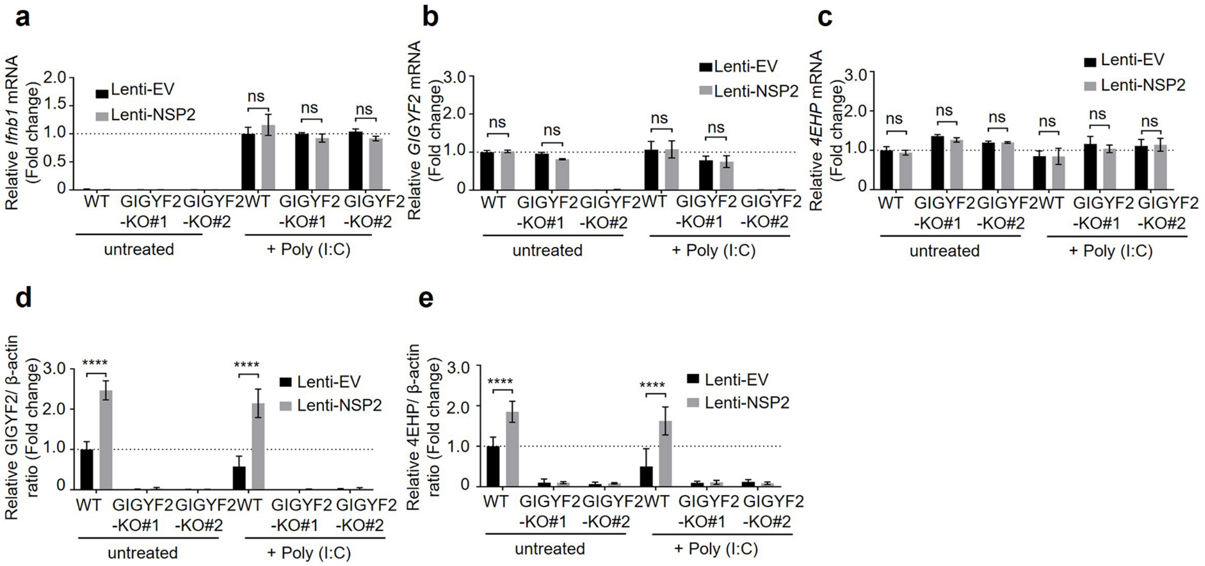

**Extended Data Figure 7. NSP2/GIGYF2/4EHP repressor complex does not affect *Ifnb1* mRNA abundance, Related to Figure 4. (a-c)** RT-qPCR analysis of *Ifnb1* (**a**)*, GIGYF2* (**b**)*, and 4EHP* (**c**) mRNAs from samples described in Fig. 4c. **(d-e)** Quantitation of expression of endogenous GIGYF2 (**d**) and 4EHP (**e**) proteins from samples described in Fig. 4d. Data are presented as mean ± SD (n=3). ns=non-significant, two-way ANOVA with Bonferroni's post hoc test.

**Extended Data Figure. 8**

**a**

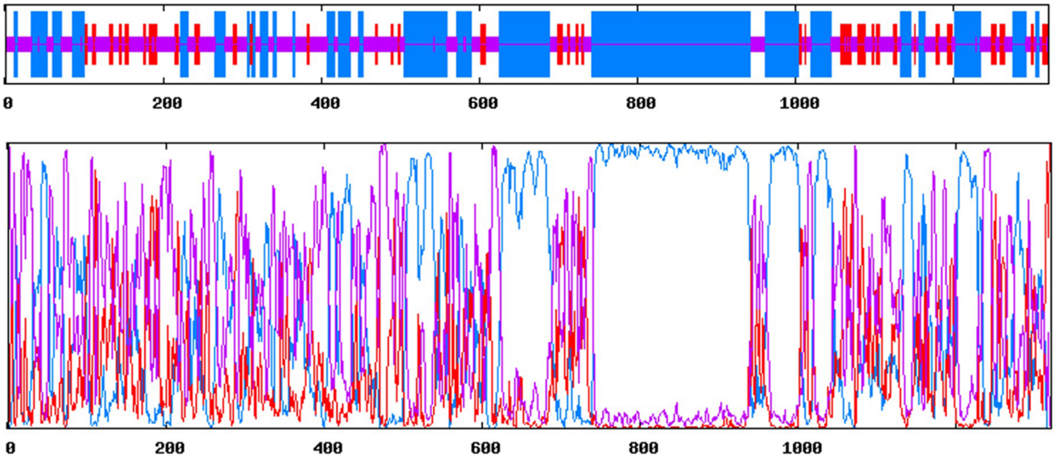


**b**

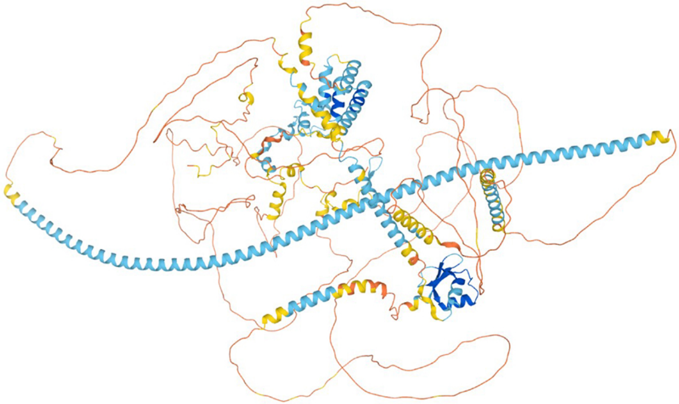

**c**

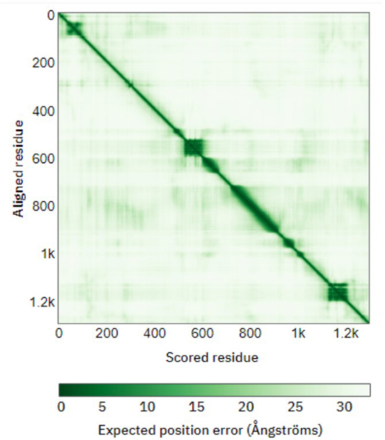

**Extended Data Figure 8. *In silico* structural analyses of human GIGYF2, Related to Figure 5. (a)** Secondary structure prediction of GIGYF2 by the Garnier-Osguthorpe-Robson IV (GOR4) method. Top is a schematic of predicted secondary structures along the primary structure of GIGYF2. Blue – regions predicted to fold into alpha helices, red – regions predicted to fold into beta sheets, purple – regions predicted to be disordered. Bottom is a graph of the probability of each residue along the primary structure of GIGYF2 to be folded into a particular secondary structure type. Blue - plot of the probability of alpha helical folding, red – plot of the probability of beta strand folding, purple – plot of the probability of disorder. **(b)** Predicted three-dimensional model of GIGYF2 from the AlphaFold Protein Structure Database. Model is coloured according to the predicted local distance difference test (pLDDT) score per residue. Dark blue – pLDDT > 90. Light blue – 90 > pLDDT > 70. Yellow – 70 > pLDDT > 50. Orange – pLDDT < 50. **(c)** Predicted alignment error (PAE) plot for the model of GIGYF2 from (b).

**Extended Data Figure. 9**

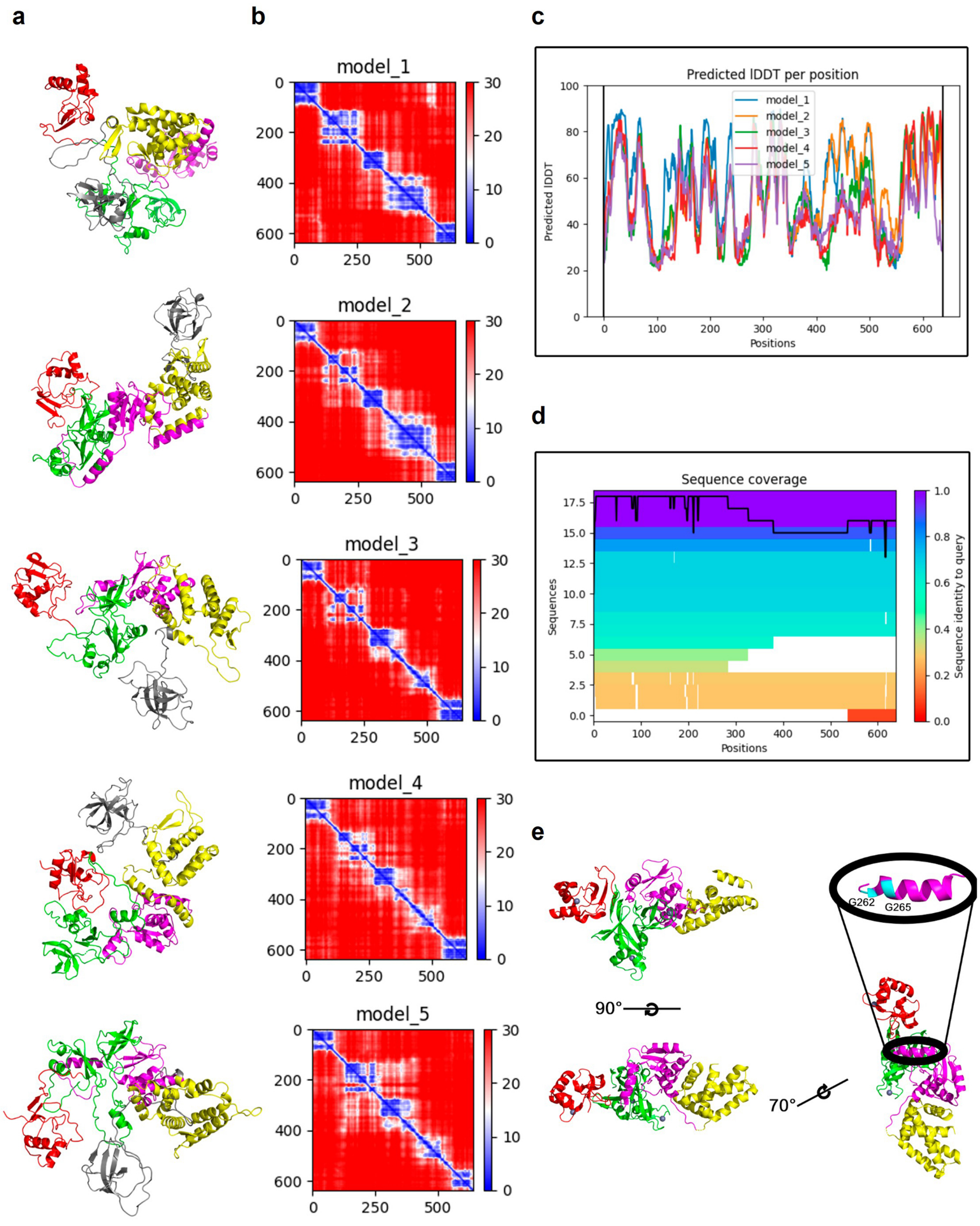

**Extended Data Figure 9. *In silico* structural analyses of SARS-CoV-2 NSP2 by AlphaFold 2, Related to Figure 5. (a)** Predicted three-dimensional models of NSP2 by AlphaFold2. Five different models for the structure of NSP2 were predicted by analyzing the full amino acid sequence of NSP2 with ColabFold. **(b)** PAE graphs for the NSP2 models from (a). **(c)** pLDDT scores per residue plots for the NSP2 models from (a). **(d)** Graph of the quality of the multiple sequence alignment per residue used as input for the prediction of NSP2 models from (a). **(e)** Position of residues G262 and G265 in the cryo-EM structure of NSP2. Annotation was done in PyMOL: red – NSP2-R1, green – NSP2-R2, magenta – NSP2-R3, yellow – NSP2-R4, cyan – G262 and G265.

**Extended Data Figure. 10**

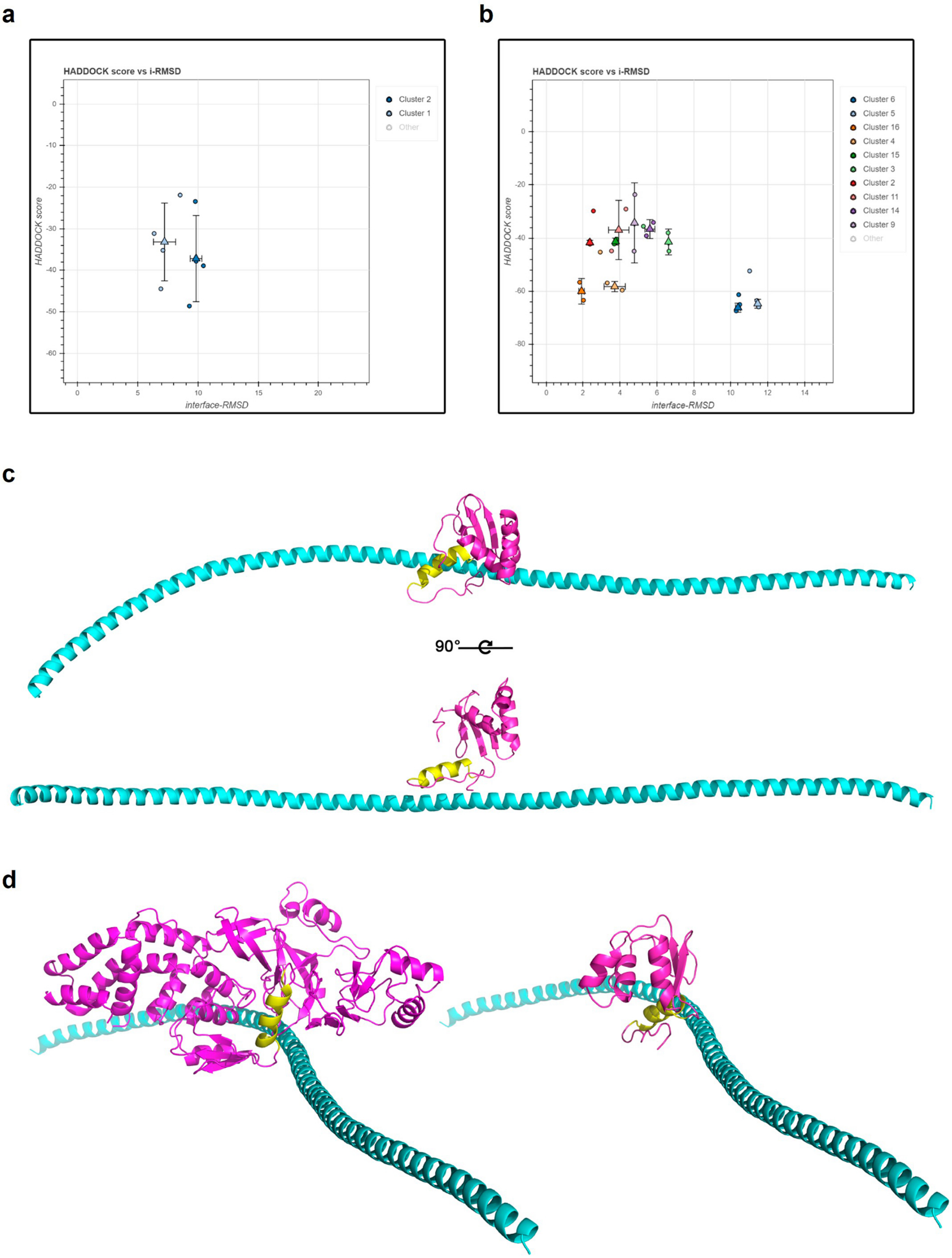

**Extended Data Figure 10. Cluster analyses of the protein-protein docking simulations between NSP2 and GIGYF2-LHR by HADDOCK 2.4, Related to Figure 5. (a)** Cluster analysis plot of the docking simulation between GIGYF2-LHR and NSP2-R3. HADDOCK score is a composite measure of the free energy of the predicted model which considers van der Waals, electrostatics and desolvation energies. Interface root mean squared distance (i-RMSD) compares the spatial similarity of the protein-protein interfacing regions between models grouped in the same cluster. Circular points – one predicted model within a cluster. Triangular points – point of mean HADDOCK score and i-RMSD within a cluster presented as mean ± 1 S.D. **(b)** Cluster analysis plot of the docking simulation between GIGYF2-LHR and the cryo-EM structure of NSP2. Same legend as (A). **(c)** Model of the interaction between GIGYF2-LHR and NSP2-R3 by HADDOCK 2.4. Annotation was done in PyMOL: magenta – NSP2-R3, cyan – GIGYF2-LHR, yellow – alpha helix possessing G262 and G265. **(d)** Comparison of the best models from both HADDOCK 2.4 docking simulations. Left model is of the docking between GIGYF2-LHR and the cryo-EM structure of NSP2. Right model is of the docking between GIGYF2-LHR and NSP2-R3 aligned to the model to the right. Annotation was performed the same as in (c) in PyMOL.

#### Extended Data Table. 1

| Name | Sequence | Application |
|---|---|---|
| IFN-β-Fwd | 5'-AAACTCATGAGCAGTCTGCA-3' | RT-PCR |
| IFN-β-Rv | 5'-AGGAGATCTTCAGTTTCGGAGG-3' | RT-PCR |
| GAPDH-Fwd | 5'-TGGGTGTGAACCATGAGAAG-3' | RT-PCR |
| GAPDH-Rv | 5'-ATGGACTGTGGTCATGAGTC-3' | RT-PCR |
| ISG56-Fwd | 5'-TCCCCTAAGGCAGGCTGTC-3' | RT-PCR |
| ISG56-Rv | 5'-GACATGTTGGCTAGAGCTTCTTC-3' | RT-PCR |
| eIF4E2-Fwd | 5'-TCCCGTTGCTGGGAGAATCTCA-3' | RT-PCR |
| eIF4E2-Rv | 5'-GTTGCTTGGTCACTGGCAGTCT-3' | RT-PCR |
| GIGYF1-Fwd | 5'-GCTGCCGTGGAGATTCGAGAAAGC-3' | RT-PCR |
| GIGYF1-Rv | 5'-CCCGCAGCCTCAGCTCCTGGC-3' | RT-PCR |
| GIGYF2-Fwd | 5'-GGGGGACATAGCCGTGATGCCTTTTAAG-3' | RT-PCR |
| GIGYF2-Rv | 5'-GGTGGAAAGGCATGGGTCAATACATTAG-3' | RT-PCR |
| GIGYF2-Fwd | 5'-GAGAAGCTGGGGAGTATTGACTGGGG-3' | RT-PCR |
| GIGYF2-Rv | 5'-CCCCTCCCCAAGAATCACTCTTCC-3' | RT-PCR |
| GIGYF2-Fwd | 5'-AACGACTGACCAGGCAGCAAGA-3' | RT-PCR |
| GIGYF2-Rv | 5'-GGAAGACAGTGCTGCTTTCTGC-3' | RT-PCR |
| Firefly luciferase-Fwd | 5'-GCCATGAAGCGCTACGCCCTGG-3' | RT-PCR |
| Firefly luciferase-Rv | 5'-TCTTGCTCACGAATACGACGGTGG-3' | RT-PCR |
| Renilla luciferase-Fwd | 5'-TCAGTGGTGGGCTCGCTGCA-3' | RT-PCR |
| Renilla luciferase-Rv | 5'-CTTTGGAAGGTTCAGCAGCTCG-3' | RT-PCR |
| GIGYF1 sgRNA#1-Fwd | 5'-CACCGTGACTACCGTTATGGGCGAG-3' | Cloning |
| GIGYF1 sgRNA#1-Rv | 5'-AAACCTCGCCCATAACGGTAGTCAC-3' | Cloning |
| GIGYF1 sgRNA#2-Fwd | 5'-CACC AGCTGGCTGACTACCGTTAT-3' | Cloning |
| GIGYF1 sgRNA#2-Rv | 5'-AAACATAACGGTAGTCAGCCAGCTC-3' | Cloning |
| GIGYF1 sgRNA#3-Fwd | 5'-CACCGAAGCTGGCTGACTACCGTTA-3' | Cloning |
| GIGYF1 sgRNA#3-Rv | 5'-AAACTAACGGTAGTCAGCCAGCTTC-3' | Cloning |
| GIGYF2 sgRNA#1-Fwd | 5'-CACCGGGATGTAATACTCCCACCAC-3' | Cloning |
| GIGYF2 sgRNA#1-Rv | 5'-AAACGTGGTGGGAGTATTACATCCC-3' | Cloning |
| GIGYF2 sgRNA#2-Fwd | 5'-CACCGAATTTATACTTCGGCAATGC-3' | Cloning |
| GIGYF2 sgRNA#2-Rv | 5'-AAACGCATTGCCGAAGTATAAATTC-3' | Cloning |
| GIGYF2 sgRNA#3-Fwd | 5'-CACCGGGAGGAACCCCTTCCACCAT-3' | Cloning |
| GIGYF2 sgRNA#3-Rv | 5'-AAACATGGTGGAAGGGGTTCCTCCC-3' | Cloning |
| 4EHP sgRNA#1 a | 5'-TGAGCTCGTGGGACGGCCGG-3' | CRISPR |
| 4EHP sgRNA#1 b | 5'-TGAAGAGATGGAAGTCACTG-3' | CRISPR |
| 4EHP sgRNA#1 c | 5'-TTGTATTCCATAATGGTGTT-3' | CRISPR |
| 4EHP sgRNA#2 a | 5'-AAAAGTGTAGTTGTACTGCA-3' | CRISPR |
| 4EHP sgRNA#2 b | 5'-TTGTATTCCATAATGGTGTT-3' | CRISPR |
| 4EHP sgRNA#2 c | 5'-GGATATATTAATGGTTTCTTTTGG-3' | CRISPR |
| GIGYF2 sgRNA#1 a | 5'-GGCAATGCTGGAGAAAGAGG-3' | CRISPR |
| GIGYF2 sgRNA#1 b | 5'-AATACGGAAAAGAATGGCAG-3' | CRISPR |
| GIGYF2 sgRNA#1 c | 5'-TGTCTTGGCCTGGTCGGAGGCATC-3' | CRISPR |
| GIGYF2 sgRNA#2 a | 5'-GATTGGTCTGAAATACTGAA-3' | CRISPR |
| GIGYF2 sgRNA#2 b | 5'-TGATGAACGGGGTTACCGAA-3' | CRISPR |
| GIGYF2 sgRNA#2 c | 5'-CGCAGCTTAAGAAAAGTAGCCGTC-3' | CRISPR |

**Extended Data Table 1. The list of primers and CRISPR guide RNA used in this study, Related to methods section.**
